# Image modifications reduce differences in natural-image encoding by retinal ganglion cells between natural and optogenetic stimulation

**DOI:** 10.64898/2026.03.06.710066

**Authors:** Varsha Ramakrishna, Tim Gollisch

## Abstract

Optogenetics is a promising approach for restoring vision after photoreceptor degeneration. Yet, little is known about how responses to natural stimuli in optogenetically treated retinas compare to photoreceptor-driven responses and how any differences might be counteracted by adjusting the stimulation. We here therefore directly compared the encoding of natural images by individual retinal ganglion cells under both conditions in mouse retinas with intact photoreceptors and channelrhodopsin expression in ganglion cells. This showed that channelrhodopsin-evoked responses display reduced dynamic range, more linear encoding of receptive-field activation, and reduced sensitivity to local spatial contrast. We thus devised image modifications, combing thresholding and scaling of pixel intensities with spatial low-pass filtering, and found that applying such modified images for optogenetic stimulation restores original encoding characteristics, yielding responses more similar to the original photoreceptor-evoked activity. These findings may help optimize stimulation of optogenetically treated retinas to achieve more natural vision in future therapeutic applications.

## Introduction

Blindness from retinal degeneration caused by diseases such as retinitis pigmentosa, age-related macular degeneration, or diabetic retinopathy affects millions of individuals worldwide (Hartong et al. 2006; Flaxman et al. 2017). These diseases often lead to a loss of photoreceptors, the main light-sensing cells in the retina. Approaches of gene replacement therapy (Bainbridge et al. 2008; Boye et al. 2013; Sahel, Roska 2013; Dalkara et al. 2016) aim at preventing or slowing down their degeneration but generally need to be developed for specific subvariants of the many forms of retinal degeneration. When photoreceptors have mostly degenerated leading to complete blindness, current therapy concepts are targeted at vision restoration through photoreceptor or stem-cell transplantation (Li et al. 2013; Singh et al. 2013; Pearson 2014; Zhao et al. 2017; Llonch et al. 2018; Temple 2023; Zerti et al. 2025), electrode-based retinal prostheses (Zrenner 2002; Nirenberg, Pandarinath 2012; Chuang et al. 2014; Palanker 2023; Holz et al. 2026), application of synthetic photosensitive molecules (Polosukhina et al. 2012; Berry et al. 2015; Tochitsky et al. 2018), or optogenetics (Lüscher et al. 2025). The latter technique is based on the expression of light-sensitive ion channels in surviving non-photoreceptor retinal neurons to convert them into artificial photoreceptors (Duebel et al. 2015; Kleinlogel et al. 2020; Simon et al. 2020; Lindner et al. 2022; Yan et al. 2023; Lüscher et al. 2025). The applied light-sensitive ion channels are typically channelrhodopsins, such as the naturally occurring Channelrhodopsin-2 (ChR2) and its genetically engineered variants.

Previous studies have shown that stimulation of retinas expressing channelrhodopsins in ganglion, bipolar, or amacrine cells of the retina can trigger neuronal light responses and restore simple light-driven behaviors in blind mice (Bi et al. 2006; Lagali et al. 2008; Lin et al. 2008; Tomita et al. 2010; Doroudchi et al. 2011; Gaub et al. 2014; Macé et al. 2015; van Wyk et al. 2015; Barrett et al. 2016; Ferrari et al. 2020; Khabou et al. 2023). Channelrhodopsin-driven responses in the retina have also been demonstrated in optogenetically treated primate retinas (Sengupta et al. 2016; Murphy et al. 2025), and first clinical trials with retinitis pigmentosa patients have reported partial recovery of vision (Sahel et al. 2021; Lam et al. 2025; Mohanty et al. 2025).

Most previous preclinical studies of retinal responses to light stimulation of channelrhodopsin in retinal ganglion cells have focused on sensitivity to flashes of light and occasional investigations of spatial resolution (McGregor et al. 2020; Gauvain et al. 2021; Lindner et al. 2021; Reh et al. 2021). Yet, improving optogenetic retinal stimulation for future vision restoration therapies requires a better understanding of optogenetically evoked responses to complex and natural visual stimuli. Given that a large part of the retinal circuitry is skipped when ganglion cells are stimulated directly with channelrhodopsin, responses to natural stimuli likely differ from normal, photoreceptor-evoked responses. To date, however, these differences have not been well characterized. Once established, such characterizations of response differences to natural stimuli might then be used to adjust the stimulation patterns in the case of optogenetic vision restoration in order to render responses more similar to normal responses for a given natural scene. Comparing normal and optogenetically evoked responses, however, is complicated by the fact that variability of responses to natural stimuli among ganglion cells is high even under wild-type conditions because different ganglion cell types respond to different visual features and each individual cell reacts according to the stimulus part within its own receptive field. Ideally, the comparison thus takes cell identity and location into account.

In the present work, we therefore assessed normal, photoreceptor-driven as well as optogenetically-driven responses in the retina on a cell-by-cell basis. To do this, we performed multi-electrode array recordings of spiking activity from retinal ganglion cells (RGCs) of a non-blind transgenic mouse line that expresses ChR2 in the ganglion cells and contains intact photoreceptors as well. This enabled us to directly compare responses of the same cells when stimulated via photoreceptors below the threshold of ChR2 activation and via ChR2 when photoreceptor signals were pharmacologically blocked. We demonstrate characteristic differences between the two scenarios, including increased receptive-field size under optogenetic stimulation, as well as reduced thresholding, reduced nonlinearities of contrast encoding, reduced spatial-contrast sensitivity, and reduced response gain in the encoding of flashed natural images. We furthermore show how these observed differences between photoreceptor-driven and ChR2-driven responses can be counteracted by modifying the natural images for optogenetic stimulation.

## Results

Improving the encoding of visual stimuli by optogenetic therapies will benefit from understanding the differences between photoreceptor-driven and optogenetically evoked neuronal activity in the retina. We therefore aimed at comparing the responses from the same retinal ganglion cells (RGCs) in these two conditions and at using the gained insight to guide modifications of the stimulation patterns in the context of optogenetics to make responses to visual stimuli more similar to those under photoreceptor stimulation. To do this, we used retinas from transgenic mice, which had intact photoreceptors as well as expression of Channelrhodopsin-2 (ChR2) in RGCs. The mice were obtained by mating animals from the Ai32 line, which carry Cre-inducible ChR2, with animals expressing Cre recombinase under the control of the VGLUT2 promoter. Staining for EYFP, the reporter that is co-expressed with ChR2, verified expression in virtually all ganglion cells of these retinas (Fig. 1A).

**Figure 1.**
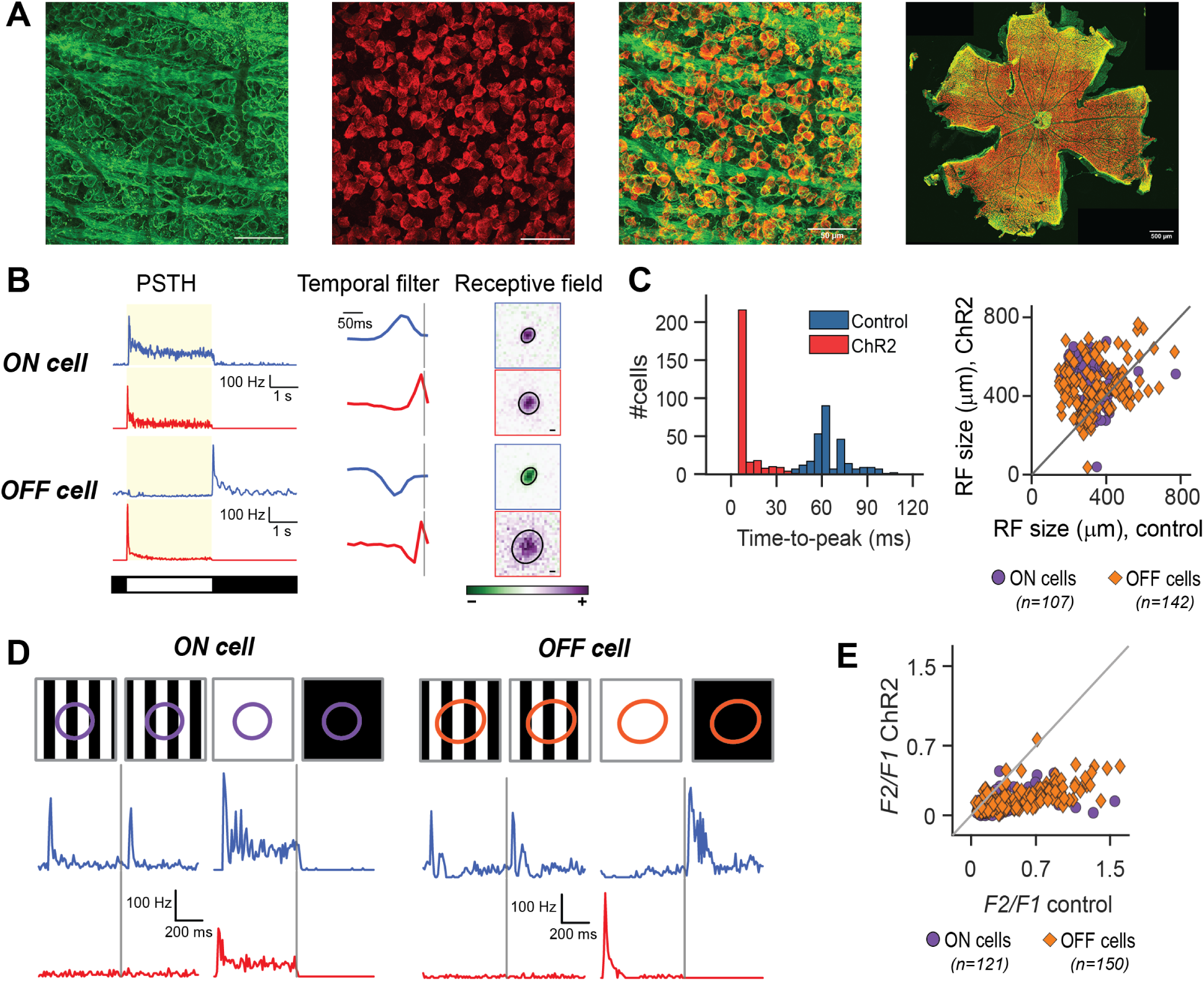
Basic characterization of ChR2-driven responses. **A.** Images of a sample retina expressing ChR2 (green, identified by staining for EYFP, which is co-expressed with ChR2) in retinal ganglion cells (red, identified by staining for RBPMS). Scale bars: 50 µm for first three images, 500 µm for rightmost image. **B.** Sample PSTHs for light-intensity steps (left), temporal filters (middle), and receptive fields (right) for an ON (top) and an OFF cell (bottom) in control (blue) and ChR2 (red) conditions. The light-on phase on the left is depicted by the shaded region. The gray vertical lines mark the time of spike for the temporal filters. Scale bar for receptive fields: 100 µm. **C.** Left: Histogram showing time-to-peak of the temporal filters for all cells in control (blue) and ChR2 (red) conditions. Right: Receptive field diameters in control versus ChR2 condition for ON cells (purple) and OFF cells (orange). **D.** PSTHs of an ON (left) and an OFF cell (right) for contrast-reversing gratings (fine stripes of 50 µm as well as broad stripes of 800 µm) under control (blue) and ChR2 (red) conditions. Top row shows the cells’ receptive-field outlines from the control condition overlaid on part of the stimulus for illustration. **E.** Comparison of nonlinearity indices *F* 2*/F* 1 from contrast-reversing gratings in ChR2 and control conditions for ON cells (purple) and OFF cells (orange).

To record spiking activity of RGCs, isolated retinas were placed on planar multi-electrode arrays. Measurements started with light intensities in the low photopic range (spanning the UV, blue, and green parts of the spectrum) for characterizing photoreceptor-driven responses (in the following referred to as control condition). In this condition, the light intensities were far below the level needed to adequately activate ChR2 so that responses can be considered as coming from a normal, wild-type retina. Subsequently, ChR2-driven spiking responses from the same RGCs were recorded by using high-intensity blue light and blocking excitatory signals in the retina with a cocktail of pharmacological drugs to avoid contributions from photoreceptors (in the following referred to as ChR2 condition). This allowed us to compare responses of individual RGCs to the same stimuli (apart from the change in light intensity and spectrum) under the two conditions.

### General characterization of ChR2-driven responses

To provide a background for the analysis of responses to natural images, we first aimed at characterizing basic response properties, such as receptive fields, temporal filters, and spatial stimulus integration, and compare these between the two stimulus conditions. We therefore started by assessing responses to simple, artificial stimuli that are commonly used for general characterizations of RGCs. For full-field step-like changes in light intensity, ChR2 was able to drive strong and reliable responses (Fig. 1B, left). ON cells responded strongly to the light-on phase in both stimulus conditions, whereas for OFF cells, as expected, response peaks switched from the light-off stimulus phase in the control condition to the light-on phase in the ChR2 condition, since the ChR2 in the cells leads to excitation by light.

Responses to spatiotemporal white noise were used to derive temporal filters and spatial receptive fields (RFs; Fig. 1B, right) under each condition by calculating spatiotemporal spike-triggered averages (STAs) separately for photoreceptor-and ChR2-driven responses. Temporal filters were found to have faster kinetics in the ChR2 condition with earlier peaks as compared to the control condition (Fig. 1C, left, time to peak 11.5 ms *±* 7.1 ms, mean *±* standard deviation, for the ChR2 condition and 66 ms *±* 13 ms for control; n = 282). This is expected, given the ChR2 opening kinetics in the range of few milliseconds (Nagel et al. 2005) and the fact that the light directly activates the ganglion cells in the ChR2 condition, bypassing the retinal circuitry. Overall, the temporal filters in the ChR2 condition are similar across cells, likely partly reflecting ChR2 channel kinetics (Nagel et al. 2005; Lin et al. 2009; Berndt et al. 2011). For example, the vast majority of cells had a time-to-peak close to 8 ms of the temporal filter in the ChR2 condition (Fig. 1C, left), in line with the opening kinetics of ChR2 at light onset (Nagel et al. 2005; Gauvain et al. 2021). Yet, some differences in temporal filter shapes remain, in particular with respect to the depth of the secondary, negative peak of the filter.

Differences between the control and ChR2 conditions were also apparent in the receptive-field sizes (obtained via a Gaussian fit to the spatial component of the STA under spatiotemporal white noise). In the ChR2 condition, RFs were larger as compared to the control condition (average RF diameter from the 1.5-*σ* contour of the Gaussian fit in the ChR2 condition: 415 µm *±* 117 µm, mean *±* standard deviation, versus 320 µm *±* 97 µm in control, *p <* 10^−18^, Wilcoxon signed-rank test, n=249, Fig. 1C, right). Viewing sample receptive fields as the ones in Fig. 1B, right, suggests that the larger receptive fields in the ChR2 condition may result from diffuse extensions beyond the original boundaries of the receptive-field center. A potential explanation is that the canonical surround inhibition is suppressed in the ChR2 condition, as bipolar-cell signals are pharmacologically blocked and amacrine cells therefore receive no light-driven inputs. Gap junction couplings between ganglion cells (Hidaka et al. 2004; Schubert et al. 2005; Trenholm, Awatramani 2019), on the other hand, should remain functional and could thereby also contribute to enlarging receptive fields.

To test integration of visual signals within the receptive-field center of RGCs in the ChR2 condition, we stimulated the retina with contrast-reversing gratings at different spatial frequencies and phases. In the control condition, many RGCs displayed responses with frequency doubling to reversals of fine gratings (50 µm bar width), indicating nonlinear spatial integration (Enroth-Cugell, Robson 1966), in line with previous studies in the mouse retina (Schwartz et al. 2012; Karamanlis, Gollisch 2021). In the ChR2 condition, however, the cells generally lost their responses to fine gratings as well as the frequency doubling and only responded to one reversal direction of coarser gratings or of full-field contrast (Fig. 1D). This indicated that bright and dark regions of fine spatial gratings could now cancel out, resulting in no net activation of the cell, as is the case for linear spatial integration.

To quantify this effect, we computed a nonlinearity index by comparing the amplitude of the maximum frequency-doubled response component to the amplitude of the coarse-grating response (see Methods). Taking the ratio of these two response components yielded a nonlinearity index, with high and low index values indicating nonlinear and linear spatial integration, respectively. We found that nonlinearity indices in the control condition covered a wide range from zero to above unity, but were generally much smaller and typically below 0.4 in the ChR2 condition (Fig. 1E; *p <* 10^−40^, Wilcoxon signed-rank test, n = 271). This shows that ChR2-driven responses in RGCs lack nonlinear spatial integration and are less sensitive to high-frequency spatial contrast.

### Responses to natural images

In order to study ChR2-driven responses to natural images, RGC spiking activity was recorded in response to photographic pictures, taken from standard databases (see Methods), in both control and ChR2 conditions. Images were presented to the retina for 200 ms each, separated by 600 ms of background illumination at mean light intensity of the images. We observed strong responses to natural images in both conditions, showing that ChR2 can drive RGCs under stimulation with natural spatial stimulus statistics (Fig. 2A).

**Figure 2.**
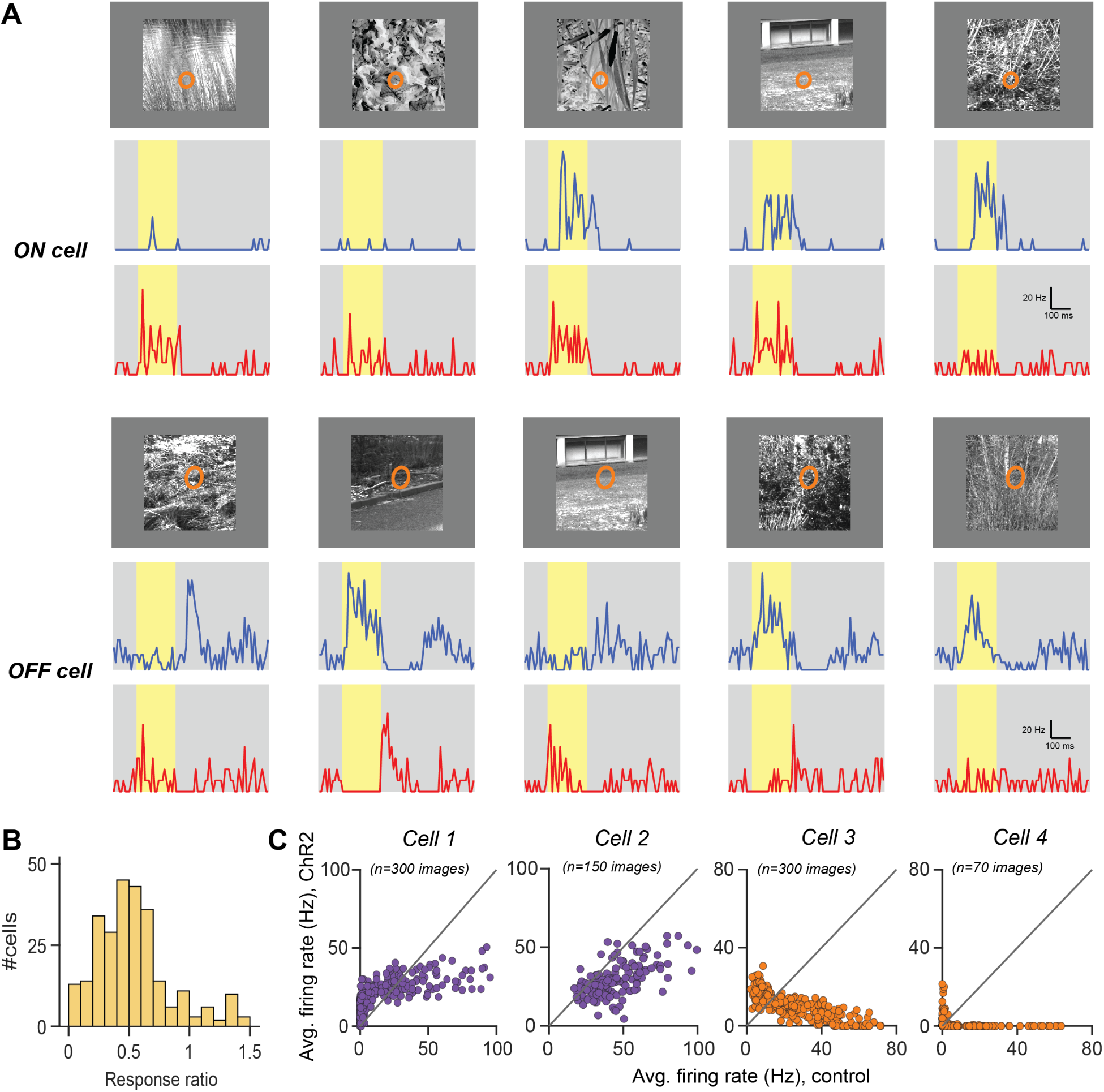
Responses of RGCs to natural images in control and ChR2 conditions. **A.** Sample images with RF outlines from the ChR2 condition and corresponding PSTHs in control (blue) and ChR2 (red) conditions for a sample ON cell (top three rows) and a sample OFF cells (bottom three rows). Brightly shaded regions in the PSTHs correspond to the time of image display and the gray regions to background illumination. **B.** Histogram showing the distribution of the response ratio over all cells (n=282), i.e., of the ratio of average responses over all images between ChR2 and control condition. **C.** Comparison of average firing rates during image flashes in control versus ChR2 condition for two ON cells (*Cell 1* and *Cell 2*) and two OFF cells (*Cell 3* and *Cell 4*). Each data point corresponds to one image, and the gray line is the identity line. *Cell 1* and *Cell 3* are the same cells as in **A**.

To assess the overall response strength under ChR2 stimulation with natural images, we averaged the number of spikes during the image presentation over all images and trials for each cell in each of the two conditions and compared this average response strength between the conditions by dividing the value from the ChR2 condition by the corresponding control value. This response ratio (Fig. 2B) was typically smaller than unity (0.63 *±* 0.51, mean *±* standard deviation over cells, *p <* 10^−28^, Wilcoxon signed-rank test, n = 282), indicating that responses were systematically smaller on average in the ChR2 condition as compared to control, yet of a similar order of magnitude. In addition, responses had shorter latency in the ChR2 condition as compared to control due to the fast channel kinetics and direct stimulation of RGCs. Furthermore, OFF cells displayed the expected inverted responses: images with strong responses at the onset in the control condition generally led to offset responses in the ChR2 condition and vice versa (Fig. 2A, bottom).

Beyond these straightforward differences between image responses in the two conditions, we observed that responses were similar between the two conditions for some images but could also diverge considerably for others. For example, the ON cell of Fig. 2A, top, responded strongly to the leftmost image in the ChR2 condition, but had essentially no response for the same image in the control condition, whereas the rightmost image yielded the opposite pattern. This demonstrates that the encoding of natural images can differ substantially between photoreceptor-driven retinal processing and direct optogenetic activation of RGCs.

For a more detailed analysis on an image-by-image basis, we calculated average firing rates for each image over the duration of the image flash and compared them for each cell between the two conditions (Fig. 2C). These comparisons revealed characteristic relationships between the image responses in the two conditions. Overall, maximal responses were typically smaller in the ChR2 condition for all cells (34 Hz *±* 16 Hz, mean *±* standard deviation, as compared to 71 Hz *±* 30 Hz in control), indicating a reduced dynamic range of the optogenetically evoked responses.

Furthermore, for most ON cells, images with stronger responses in the control condition also tended to generate stronger responses in the ChR2 condition, yet this relationship was neither strict (rank-order correlations, measured by Spearman’s *ρ*, were on average 0.69 *±* 0.14, mean *±* standard deviation, n = 125 ON cells) nor linear. Typically, data points lay in concavely bent region (see *Cell 1* in Fig. 2C for an example); weakly activating images yielded relatively stronger responses in the ChR2 condition, but strongly activating images generated stronger responses in the control condition. Some ON cells also displayed a more linear relationship between responses in the two conditions (see *Cell 2* in Fig. 2C for an example), especially when baseline activity led to a lack of images with near zero firing rate.

For OFF cells (e.g., *Cell 3* and *Cell 4* in Fig. 2C), the cloud of data points displayed an inverted shape as compared to ON cells, reflecting the switch from OFF-type to ON-type contrast preference. Correspondingly, rank-order correlations were generally negative, though absolute values were comparable to those of ON cells (Spearman’s *ρ* = *−*0.72 *±* 0.14, mean *±* standard deviation, n = 157 OFF cells), indicating a similar match (apart from the inversion) between the two conditions of the ordering of images according to response strength.

To better understand the differences in the image-elicited responses between the two conditions, we computed image-response curves for each cell under each condition. These image-response curves relate the firing rates during the image flashes to the activation of the receptive field by the images. The receptive-field activation for each image was calculated as the weighted mean stimulus intensity, *I_mean_*, with weights given by the receptive field estimate (Fig. 3A, see also Methods).

**Figure 3.**
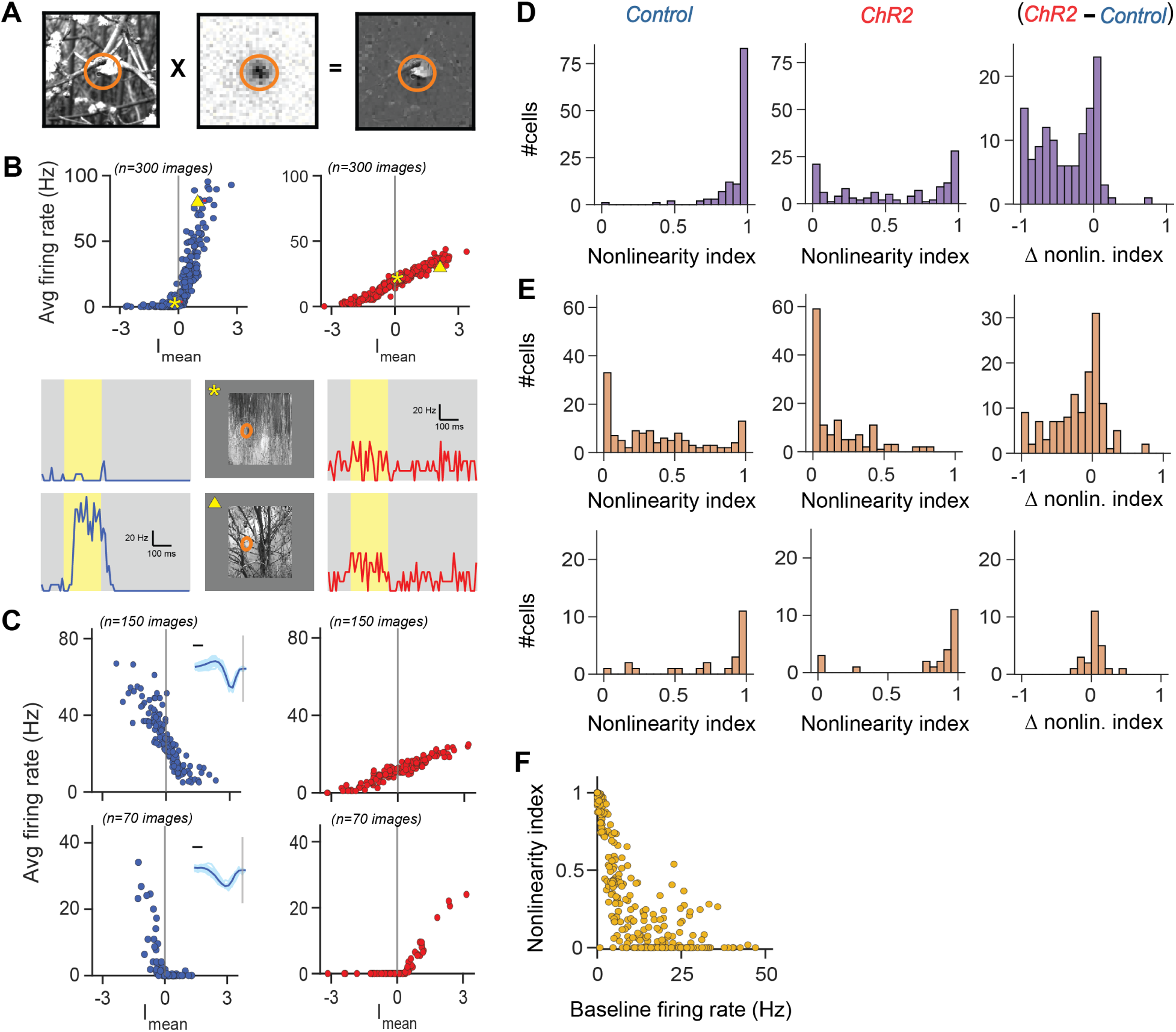
Image-response curves. **A** Schematic for calculating *I_mean_* by weighting the image pixels with the RF estimate (ellipse denoting outline) and averaging. **B.** Top: Image-response curves of a sample ON cell in control (blue) and ChR2 (red) condition. Symbols mark the data points corresponding to the examples below. Bottom: Example responses of the cell to two images in control (blue) and ChR2 (red) condition. Ellipses in the images mark the RF contours from the ChR2 condition. **C.** Image-response curves for both conditions (left/blue: control; right/red: ChR2) of a biphasic (top) and a monophasic (bottom) OFF cell. (Insets display all temporal filters of the corresponding cluster, scale bar: 50 ms). **D.** Nonlinearity indices of image-response curves for all ON cells (n=125) in control (left) and ChR2 (middle) condition and their differences Δ (right). **E.** Same as **D** for biphasic (top, n=133) and monophasic (bottom, n=24) OFF cells. **F.** Relationship between baseline firing rate and nonlinearity of image-response curves across all cells in the ChR2 condition.

The image-responses curves revealed differences between the two conditions in how the cells were activated by the images. In particular, the intensity-response relationship was generally more linear with less thresholding than in the control condition. As seen for the sample ON cell in Fig. 3B, firing rates during image presentation were often near zero for small or negative *I_mean_* values in control, whereas in the ChR2 condition, there was a continuous increase of firing rate with *I_mean_* even for negative *I_mean_*values, with substantial firing for images with *I_mean_* near zero (see responses for the sample image marked by the star symbol in Fig. 3B). As illustrated by the sample response traces below the image-response curves, the elevated firing rate for *I_mean_ ≈* 0 in the ChR2 condition is connected to an elevated background activity. This allows the firing rate to drop further below background level as *I_mean_*becomes more negative, supporting the linear relationship between receptive-field activation (*I_mean_*) and response with no or little thresholding.

In the positive range of *I_mean_*, on the other hand, firing rates in the control condition increased much more strongly than in the ChR2 condition, leading to higher firing rates for strong receptive-field activation (see responses for the sample image marked by the triangle symbol in Fig. 3B).

For OFF cells, image-response curves in the ChR2 condition had a reversed shape compared to the control condition, as expected (Fig. 3C). Beyond this, we found cells with rather linear image-response curves in both conditions (see example in Fig. 3C, top) and other cells with nonlinear, thresholded curves (Fig. 3C, bottom).

A quantitative analysis of the shape of the image-response curves corroborated these observations. To assess the degree of nonlinearity of the image-response curves, we compared the average slopes of the curves in the left and right half. Concretely, we calculated a nonlinearity index by fitting straight lines to the two parts of an image-response curve with *I_mean_ >* 0 and *I_mean_ <* 0, respectively, and taking the ratio of the difference and sum of the two obtained slope values (see Methods). Index values near unity and zero thus indicate nonlinear and linear curves, respectively.

This showed that ON cells in our dataset generally had highly nonlinear image-response curves in the control condition and strongly reduced nonlinearity indices in the ChR2 condition, with many image-response curves becoming fairly linear (Fig. 3D). Interestingly, for OFF cells, we found that the degree of nonlinearity of the image-response curves was related to the shape of the temporal filter of the cell, as assessed under white-noise stimulation in control condition. After separating OFF cells into those with monophasic filters (n = 24) and those with biphasic filters (n = 133), based on a principal-component analysis and K-means clustering (see insets in Fig. 3C for the filters of the two clusters), we found that biphasic OFF cells generally had more linear image-response curves, in particular in the ChR2 condition, whereas monophasic OFF cells responded nonlinearly to natural images in both conditions (Fig. 3E, see also examples of Fig. 3C).

Furthermore, for biphasic OFF cells, nonlinearity indices generally decreased when switching to the ChR2 condition, as seen by the negative difference between indices from the ChR2 and control condition (Fig. 3E, top-right), indicating that the biphasic OFF cells responded more linearly in the ChR2 condition as compared to control (Fig. 3E). By contrast, the monophasic cells had high nonlinearity indices in both the conditions with little change, as seen by the index differences, which cluster around zero (Fig. 3E). Thus, unlike ON cells and biphasic OFF cells, the monophasic OFF cells kept their nonlinear encoding of receptive field activation by natural images, This suggests that the degree of nonlinearity of image encoding by OFF cells is at least partly a cell-intrinsic property not inherited from presynaptic circuitry and thus indicates a cell-type-specific contribution to image processing by ChR2 in ganglion cells.

Overall, a primary difference in the encoding of natural images between the control and the ChR2 condition seems to be that the nonlinearity of the image-response curves in the control condition, stemming from thresholding near *I_mean_* = 0, is lost or reduced in the ChR2 condition because firing rates keep decreasing as *I_mean_*becomes more negative. This requires the firing rate at *I_mean_* = 0 to be substantially above zero, and the simplest explanation for this is a corresponding baseline firing rate that occurs also in the absence of a stimulus, as in the example of Fig. 3A. Indeed, when comparing the nonlinearity index to the baseline firing rate in the ChR2 condition, we found an inverse relationship (Spearman’s *ρ* = *−*0.86), so that a small nonlinearity index was typically accompanied by elevated baseline firing, whereas nonlinearly responding cells generally had lower baseline firing rates (Fig. 3F). Thus, the elevated baseline firing rate under ChR2 stimulation and the effects on the encoding natural images are an important aspect to consider when translating visual scenes in optogenetic stimulation for therapeutic applications.

### Spatial contrast sensitivity in natural images in the ChR2 condition

Natural images are rich in spatial structure coming from textures and object boundaries, and many RGCs are sensitive to such structure in the image part that falls onto the receptive field center (Schwartz et al. 2012; Turner, Rieke 2016; Karamanlis, Gollisch 2021; Liu et al. 2022; Sridhar et al. 2026). Our analyses of responses to contrast-reversing gratings (Fig. 1D,E) had indicated that the sensitivity to fine spatial structure is considerably reduced in the ChR2 condition as compared to control, at least for these artificial probe stimuli. Examining the image-response curves suggests that this difference in spatial contrast sensitivity may also influence the encoding of natural images. In particular, the data points in the image-response curves often displayed a substantial vertical spread in the control condition, indicating that images with similar *I_mean_* can elicit different responses (Fig. 4A, top, for example, see the marked responses corresponding to the two sample images). Thus, other features beside mean light intensity in the RF, such as spatial structure in the image, likely influence the responses. Image-response curves for the ChR2 condition, on the other hand, were often much “tighter” with less spread in the vertical direction (e.g., Fig. 4A, middle), showing that mean light intensity inside the RF better reflected the response strength with less influence by other features.

**Figure 4.**
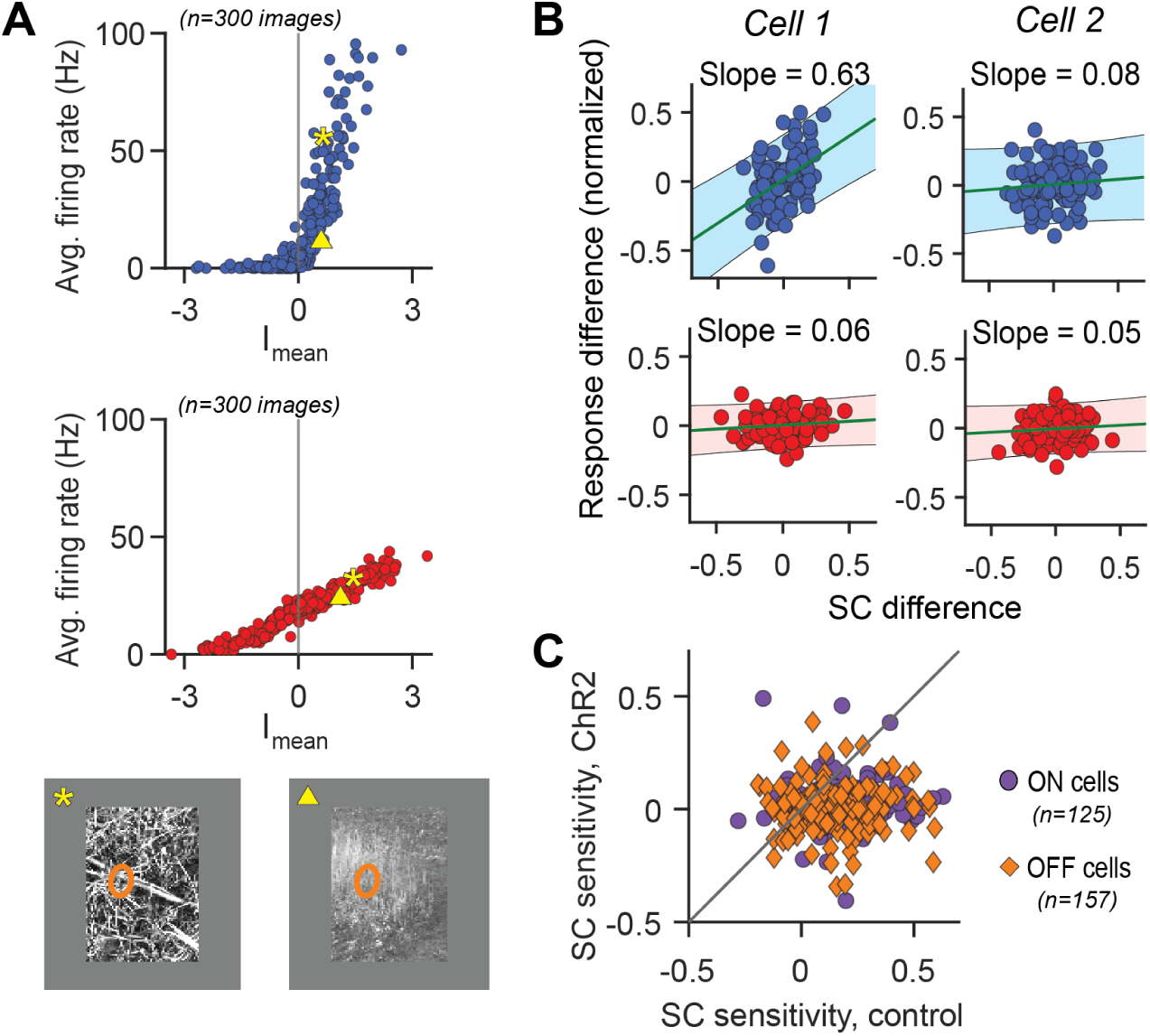
Spatial contrast sensitivity of RGCs for natural images. **A.** Image-response curves for a sample cell (same cell as in Fig. 3B) in control (blue) and ChR2 condition (red), with data points for two sample images marked by symbols. These images (shown below, with ellipses indicating the RF of the sample cell in the ChR2 condition) produced similar *I_mean_*for the sample cell but triggered markedly different responses in the control condition with the difference reduced in the ChR2 condition. **B.** Difference in local spatial contrast versus difference in response for pairs of natural images in control (blue) and ChR2 (red) conditions, for two further sample cells. Lines and shaded regions show fitted linear regression with 95% confidence intervals. The slope of the regression line is noted above each plot. **C.** Spatial contrast sensitivity in control versus ChR2 condition for ON (purple) and OFF cells (orange). The gray line marks identity.

To test this hypothesis more directly and relate the spread of data points in the image-response curves to the spatial structure in the images, we computed the spatial contrast (SC) of each image inside a cell’s receptive field center as the weighted standard deviation of pixel intensity values in the image (weights given by the RF). To relate the SC values of the images to the evoked responses in a way that is roughly independent of the mean light intensity signal *I_mean_*, we divided the images into pairs with similar *I_mean_*values (see Methods) and then computed the differences in both SC and evoked firing rate for each pair. If spatial contrast has an excitatory effect on a cell’s activity beyond what is mediated by *I_mean_*, a positive correlation between these differences is expected (Karamanlis, Gollisch 2021; Liu et al. 2022), and we therefore took the slope of a fitted regression line as the cell’s spatial contrast sensitivity (SC sensitivity) under natural images (Fig. 4B).

We observed that RGCs displayed various kinds of spatial contrast sensitivity in the control condition, from a fairly steep dependence of the response difference on the SC difference with correspondingly positive SC sensitivity values to a lack of a correlation and thus SC sensitivity near zero (Fig. 4B, top, and Fig. 4C). In the ChR2 condition, on the other hand, the SC sensitivity values decreased substantially compared to control (*p <* 10^−6^, Wilcoxon signed-rank test, n = 282) and spread around zero (Fig. 4B, bottom, and Fig. 4C), indicating no preference for spatial contrast in the images. Thus, the reduced sensitivity to spatial contrast of RGCs under direct optogenetic stimulation generalizes from reversing gratings to natural images.

### Modifying natural images to elicit more natural ChR2-driven responses

The analyses above have shown that ChR2-driven responses of RGCs differ characteristically from photoreceptor-driven responses to natural images. In particular, the cells had reduced dynamic range and reduced spatial-contrast sensitivity as well as more linear image-response curves with less thresholding in the ChR2 condition. These generic differences suggest that certain modifications of the images used for stimulation in the ChR2 condition may help in making responses generally more similar to the control condition. For example, thresholding the light intensity in the images may effectively emulate a thresholding effect of the retina and thereby make image encoding in the ChR2 condition more similar to the control condition. Alternatively, one might think that reducing overall light intensity in the ChR2 condition can create thresholding, as darker parts in the image would then fall below the activation threshold of ChR2 and background activity would be reduced. This, however, would come at the cost of overall weakening ChR2 activation and thus reducing maximal responses and dynamic range further below the level of control responses.

Instead, we here therefore selected transformations that change the spatial structure of the images while keeping the same range of applied light intensities as before. We implemented a threshold in these transformations by first subtracting the level of background illumination from the intensity of each image pixel and then clipping resulting negative pixel intensities at zero so that effectively all pixels below the previous background intensity are set to black. Subsequently, we scaled the images by a factor of two in order to restore the maximum level of pixel intensities, aiming at providing strong activation of ChR2 by brigh image parts. These thresholded and scaled images were presented to the retina in the ChR2 condition on a black background (Fig. 5A), consistent with the fact that pixel intensities corresponding to the previous background level are now set to zero.

**Figure 5.**
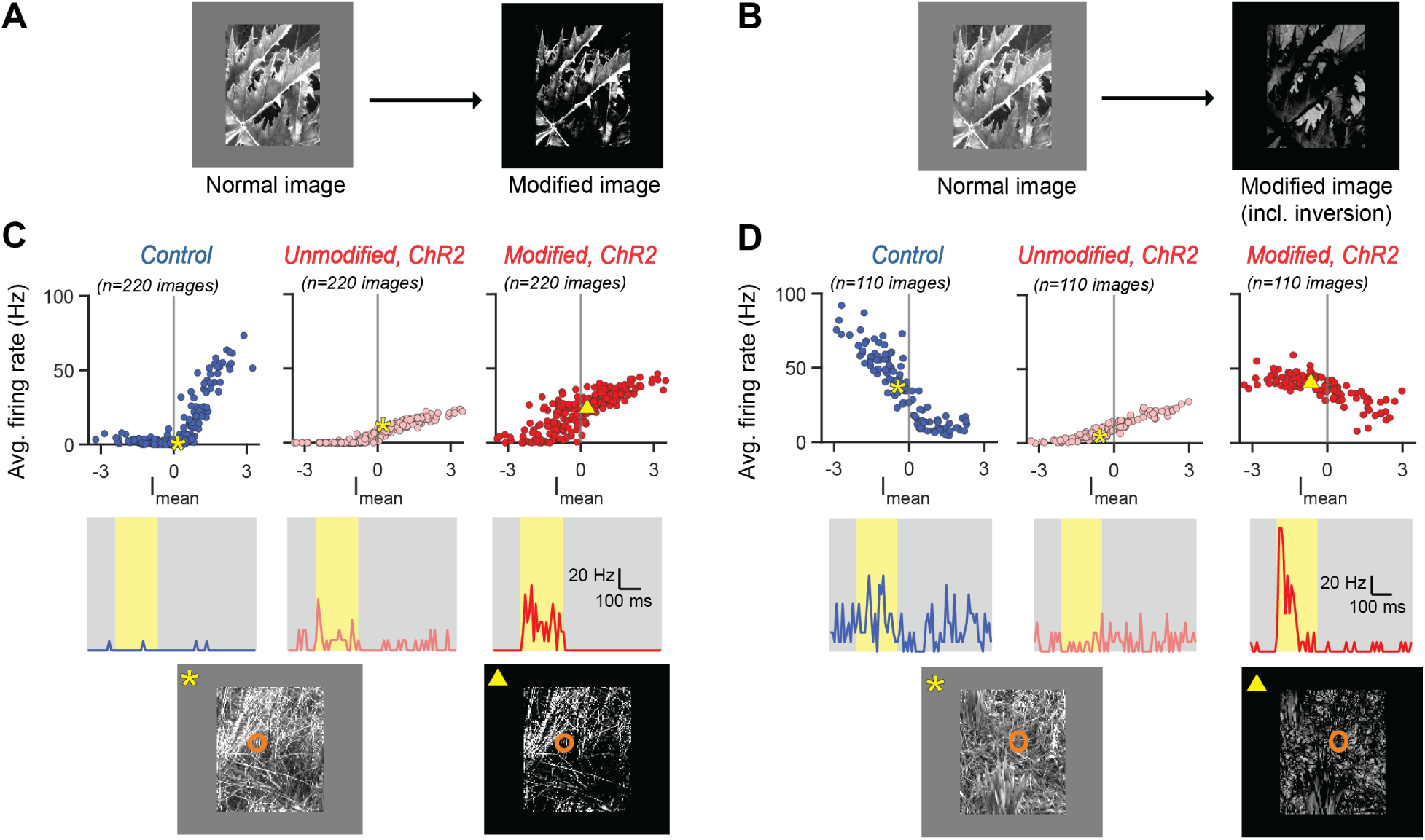
Effect of image thresholding and scaling on RGC responses. **A.** Schematic of image modifications, showing the thresholding with the subtraction of the background intensity and the scaling to match the original range of intensity values. **B.** Schematic of image modifications including the contrast inversion intended for OFF cells. **C.** Image-response curves for a sample ON cell to unmodified images in control (blue, left), unmodified images in ChR2 (pink, middle), and modified images in ChR2 (red, right) condition. Sample images with overlaid RFs from the ChR2 condition along with measured PSTHs are shown below, with symbols marking the corresponding data points in the image-response curves above. **D.** Same as **C** for a sample OFF cell.

To compute the image-response curve from the responses to the modified images, we retained the original *I_mean_*values from the unmodified images. The rationale behind this approach is that the image modification is considered part of the stimulus encoding process so that the combination of image modification and ChR2 activation of RGCs leads to responses that represent the original, unmodified image in as natural a fashion as possible.

For OFF cells, in order to make responses directly comparable to the control condition, we counteracted the switch from OFF-type to ON-type contrast preference in the ChR2 condition by additionally flipping the sign of the pixel contrast values such that dark pixels became bright and vice versa (Fig. 5B). This flipping occurred prior to the thresholding and scaling operations. As for ON cells, the *I_mean_* values for the image-response curves of OFF cells were computed from the original, non-flipped images, as the flipping is also considered part of the encoding process.

We found that these modifications led to a reduction in baseline firing rate and increased maximal responses, as well as an inversion of the direction of increasing firing rate in the image-response curves for OFF cells, as desired (Fig. 5C,D). However, we also observed an undesired increase of responses to images with low *I_mean_* values (or high *I_mean_* values for OFF cells) so that the threshold effect was not apparent in the image-response curves obtained for the modified images. By examining sample images (e.g., Fig. 5C,D, bottom), the apparent reason was readily identifiable. As the thresholding operation set all pixels below the original background intensity to the same level, even small spots of residual pixels above the background level could now lead to a net intensity above background. Given the spatial structure of natural images, almost every image thereby had a net positive activation, as no cancellation by image parts below the background level could occur.

### Including spatial blurring in the image modifications

To correct for these spurious responses triggered by small spots above background intensity in the modified images, we added further image modifications (Fig. 6A). In particular, we applied a spatial filter to the images prior to the thresholding and scaling in order to blur out small bright regions surrounded by darker image parts. The blurring lowers the intensity of small bright spots, thus reducing or suppressing the activation by bringing the spot intensity near or below the threshold at background intensity. Such spatial filtering is also part of the natural retinal signal processing before nonlinear rectification occurs, for example, by the bipolar-cell receptive fields before synaptic transmission or by the ganglion-cell receptive field before spike generation. We therefore tested two spatial scales of blurring, both implemented by convolution with a two-dimensional Gaussian filter, but using either a standard deviation *σ* of 33 µm or of 90 µm, which roughly correspond to spatial scales of bipolar or ganglion cell receptive field centers, respectively.

**Figure 6.**
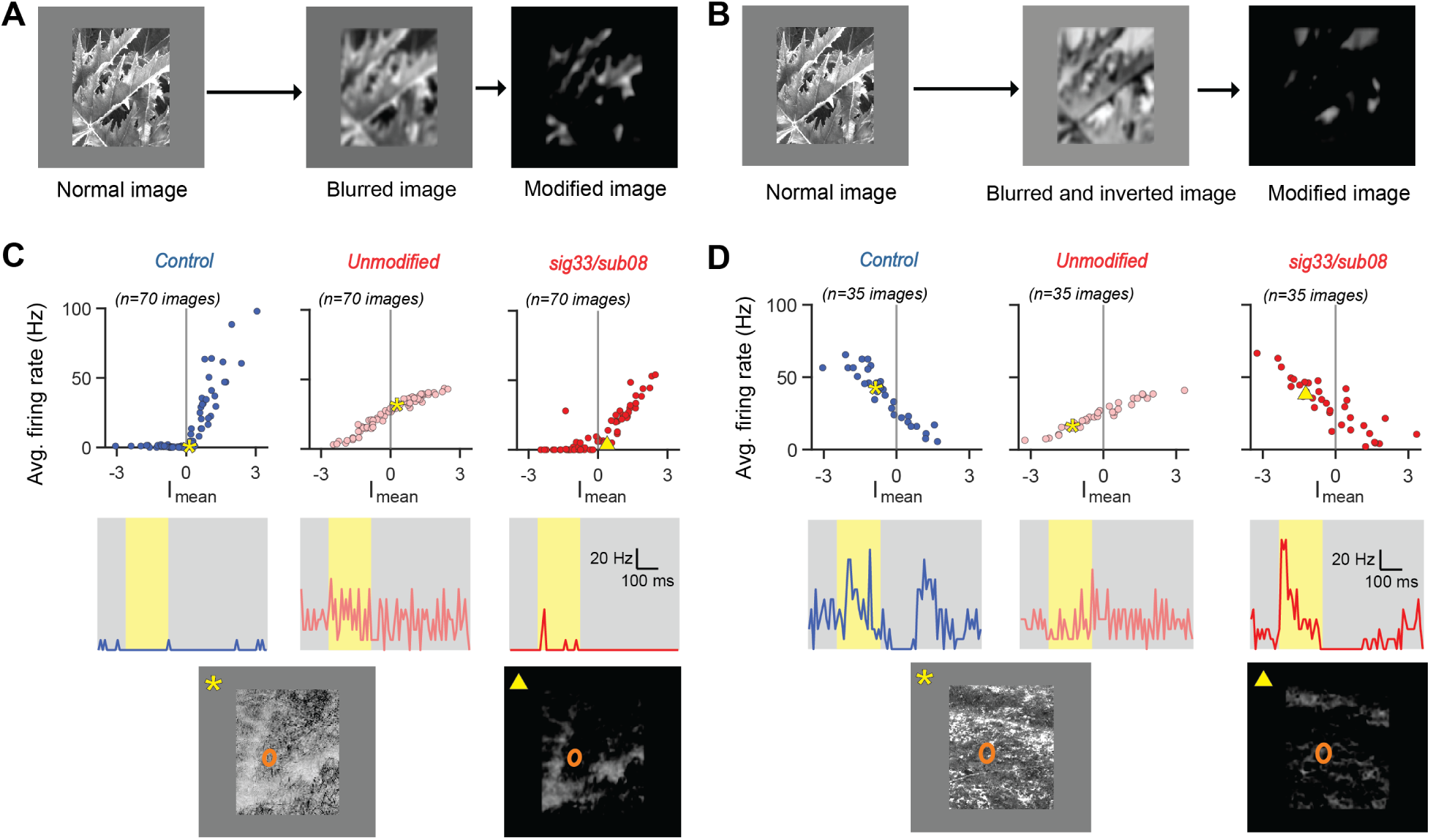
Effect of image modifications that include blurring, thresholding, and scaling. **A.** Schematic of image modifications, showing the blurring and subsequent thresholding and scaling of pixel intensities. **B.** Schematic of image modifications including the contrast inversion intended for OFF cells. **C.** Response curves for a sample ON cell to unmodified images in control (blue, left), unmodified images in ChR2 (pink, middle), and modified images (*sig33/sub08*) in ChR2 (red, right) condition. Sample images with overlaid RFs from the ChR2 condition along with measured PSTHs are shown below, with symbols marking the corresponding data points in the image-response curves above. **D.** Same as **C** for a sample OFF cell.

To further curtail overshooting activation, we also adjusted the thresholding to subtract not only the original background intensity, but also an additional 8% or 16% of the maximal possible pixel value. The subsequent scaling was adapted accordingly so as to restore the original available range of intensity values of the images. The additional subtraction was applied to help create a thresholding effect similar to the one in the control image-response curves by suppressing responses to images that trigger no or little response under control conditions (e.g., images with *I_mean_ ≈* 0).

To keep the different combinations of spatial-filtering scale and additional subtraction level apart, images modified by using, say, spatial filtering with sigma 33 µm and additional subtraction of 8% are in the following indicated as *sig33/sub08* and so on. For OFF cells, we again inverted the image contrast by flipping the sign of the pixel contrast values as described in the previous section and then applied the spatial filtering, subtraction, and scaling as described above (Fig. 6B). As before, these modified images were presented to the retina in the ChR2 condition, and the image-response

curves were computed with *I_mean_* values obtained from the unmodified images and compared across all conditions.

Figures 6C and 6D show responses for the control condition and the ChR2 condition using original images as well as images with the *sig33/sub08* modification for an ON and and OFF cell, respectively. For both cases, we see that the modification managed to recreate image-response curves similar to the control condition. In particular, responses to images that yielded little activity in the control conditions were now also suppressed in the ChR2 condition, unlike for the original images in the ChR2 condition. For the ON cell, the final image-response curve had a similar rectification as the control curve, and both cells now displayed maximal responses under ChR2 that were closer to the maximal control responses than for unmodified images.

### Comparison of image modifications with different parameters

Figure 7 shows image-response curves with unmodified images in both conditions as well as with four tested image modifications in the ChR2 condition for four sample cells. The examples demonstrate that all modifications can at least approximately restore the original nonlinear shape of the curves for the two sample ON cells (*Cell 1* and *Cell 2*), but also the milder thresholding apparent for the two OFF cells (*Cell 3* and *Cell 4*). Without image modifications, on the other hand, the image-response curves for these examples were again rather linear (second column). Furthermore, maximal responses over all images in the ChR2 condition generally reached a higher level with image modifications than without in the ChR2 condition.

**Figure 7.**
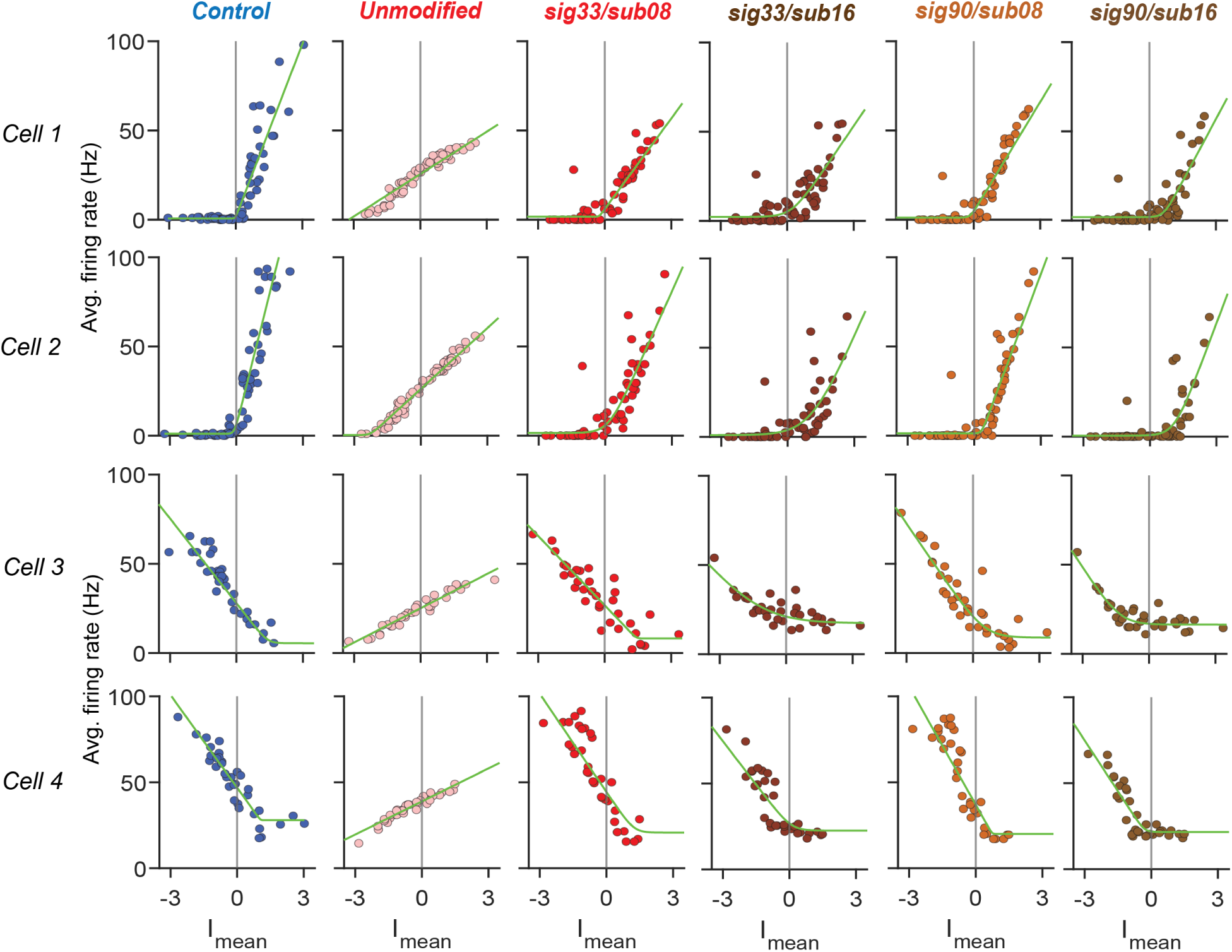
Image-response curves for the control condition (blue) and the ChR2 condition with different types of image modifications (different shades of red; type of modification indicated on top). Data are shown for two ON cells (*Cell 1* and *Cell 2*) and two OFF cells (*Cell 3* and *Cell 4*). Green lines are fitted softplus functions.

Closer inspection reveals that not all tested image modifications performed equally well. In particular, the modifications *sig33/sub16* and *sig90/sub16* display noisier curves with weaker maximal responses and a thresholding that appears too strong when compared to the control image-response curves. This indicates that for these modifications, the subtraction level was indeed too high and that the modifications *sig33/sub08* and *sig90/sub08* with the more moderate threshold appear more suitable.

In order to perform a more quantitative comparison across all cells, we first fitted a parameterized rectifying curve (softplus function, see Methods) to each image-response curve. The fitted functions managed to generally capture the structure of the image-response curves in the various conditions nicely (Fig. 7), including the varying thresholding and the different response gains. We then used the fits to compute a distance measure between the image-response curve in the control condition and the curve in the ChR2 condition with a given image modification. To do so, we subtracted the fitted control curve from the fitted ChR2 curve and used the average of the absolute difference values, averaged across the set of *I_mean_*values from the control condition data, as a distance measure. Smaller values of the distance indicate more similar image-response curves. For OFF cells, we additionally flipped the fitted curve for unmodified images in the ChR2 condition (i.e., without image inversion) across the *y*-axis before the subtraction to match the curve’s rise direction to the control condition.

We first evaluated which of the image modifications (including the case of unmodified images) resulted in the best match to the control-condition image-response curve for each cell by selecting the modification with the smallest distance. We found that two types of image modifications, *sig33/sub08* and *sig90/sub08*, stood out among the tested set as being best for a large proportion of cells (Fig. 8A).

**Figure 8.**
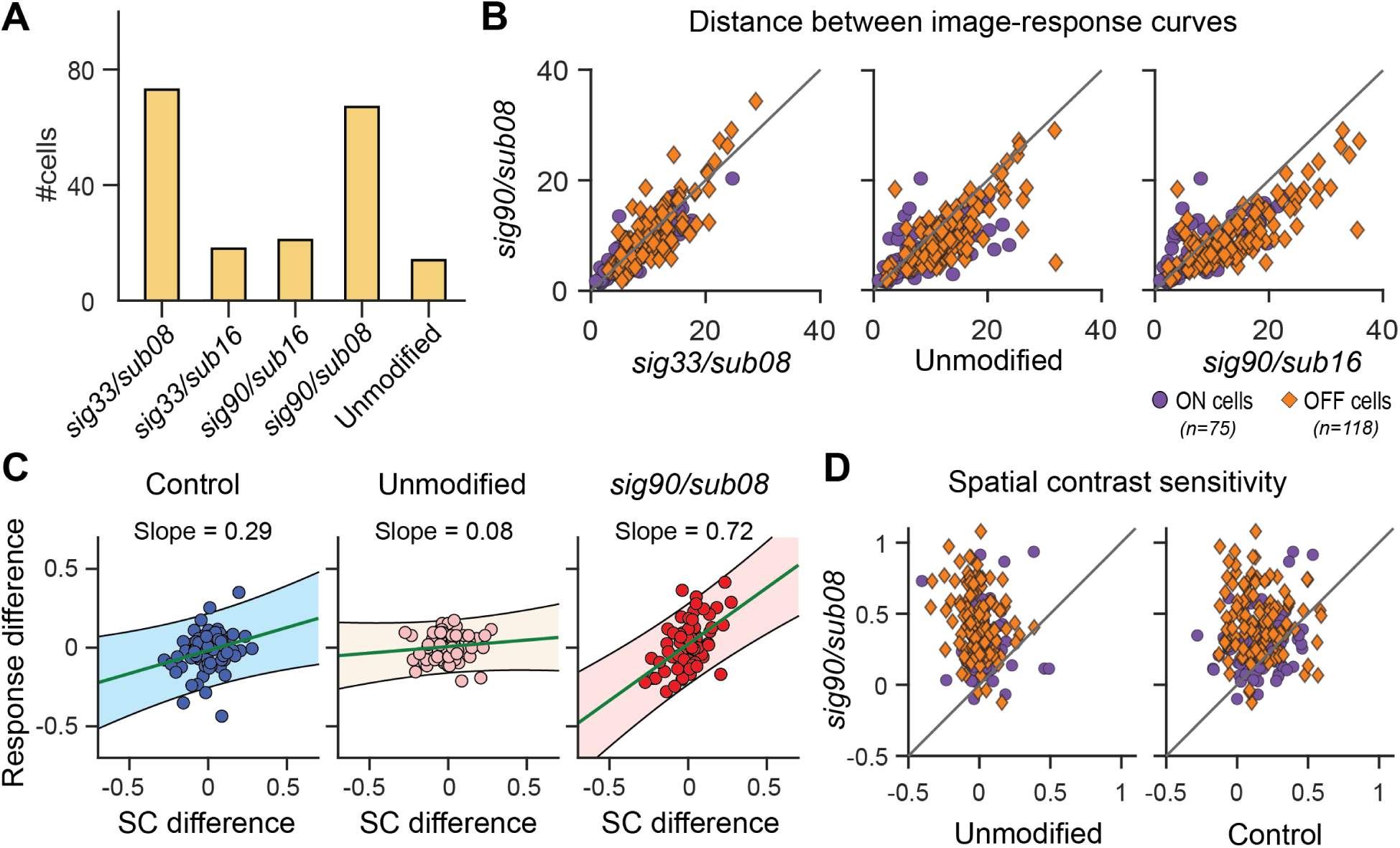
Comparison of different types of image modifications. **A.** Histogram showing for each tested modification the number of cells which had this modification as the best one (as assessed by the minimum distance between the response curves with the modified images in the ChR2 condition versus the control condition). **B.** Comparison of the distance measure for the *sig90/sub08* modification with the *sig33/sub08* modification (left), with the unmodified images (but including the inversion in the case of OFF cells; middle), and with the *sig90/sub16* modification (right) for ON (purple) and OFF (orange) cells. **C.** Difference in local spatial contrast versus difference in response (normalized to maximum response) for pairs of natural images in control condition (left, blue), ChR2 condition with unmodified images (middle, pink), and ChR2 condition with modified images (right, red) for a sample cell, shown as in Fig. 4B. Lines and shaded regions show fitted linear regression with 95% confidence intervals. The slope of the regression line is noted above each plot. **D.** Comparison of the spatial contrast sensitivity under natural images for the *sig90/sub08* modification to the spatial contrast sensitivity for unmodified images in the ChR2 condition (left) and in the control condition (right) for ON (purple) and OFF cells (orange).

Comparing the distance measures for different modifications on a cell-by-cell basis (Fig. 8B, right) showed that the performance of the *sig33/sub08* and *sig90/sub08* were generally similar, as both modifications produced similar distances of image-response curves from the control condition curves (Fig. 8B, left; *p* = 0.11 for all cells; *p* = 0.07, n=75 for ON cells; *p* = 0.39, n=118 for OFF cells; Wilcoxon signed-rank test). Compared to unmodified images, both these modifications generally resulted in improvements. The *sig90/sub08* modification, for example, led to smaller distances as compared to unmodified images for the majority of cells (Fig. 8B, middle, *p <* 10^−6^ for all cells; *p* = 0.0026, n=75 for ON cells; *p <* 10^−6^, n=118 for OFF cells; Wilcoxon signed-rank test). Yet it is noteworthy that some cells displayed similar distances or even smaller distances with no image modifications compared to *sig90/sub08*, indicating that the overall ability to match the image-response curve in the control condition depends on the particular cell at hand and that the applied image modifications are not optimal for all cell types. The modifications with the higher additional subtraction level *sig90/sub16* and *sig33/sub16*, on the other hand, generally displayed poorer performance. For example, the comparison of distance values for *sig90/sub16* with those for *sig90/sub08* showed a clear preference with smaller values for the latter (Fig. 8B, right, *p <* 10^−6^; *p* = 9.3 *×* 10^−4^, n=75 for ON cells; *p <* 10^−6^, n=118 for OFF cells; Wilcoxon signed-rank test).

The applied modifications had been designed to address specific differences in the image-response curves between the control and the ChR2 condition, in particular, the reduced nonlinear thresholding and reduced dynamic range under optogenetic stimulation with unmodified images. Another aspect of the ChR2 activation had been the loss of nonlinear spatial integration, as seen in the absence of frequency doubling for contrast-reversing gratings (Fig. 1D,E) and in the reduced spatial contrast sensitivity with unmodified natural images (Fig. 4). We therefore checked whether the applied image modifications also had an effect on spatial contrast sensitivity.

We assessed the sensitivity to spatial contrast as before (Fig. 4) by dividing the images into pairs with similar mean-intensity signals *I_mean_* in the receptive field and then relating the difference in the evoked firing-rate response to the difference in the spatial contrast for each image pair (Fig. 8C). We did this separately for responses in the control condition, in the ChR2 condition with unmodified images, and in the ChR2 condition with modified images. For the latter, we used the unmodified images to compute the *I_mean_* values for pairing images as well as to compute the local spatial contrast of the images inside the receptive fields. As for the assessment of image-response curves with modified images, the rationale is that the image modifications are considered part of the encoding process, and we are here interested in whether activity differences are informative about spatial contrast differences in the original images. Like before, the slope in the plot of response differences versus spatial-contrast differences can be used to identify spatial contrast sensitivity, with a positive value indicating high sensitivity to spatial contrast and a slope value near zero suggesting no effect of spatial contrast on the responses.

For the sample cell of Fig. 8C, the dependence of responses on spatial contrast in the control condition (as visible in the positive slope of the regression line) disappeared in the ChR2 condition for unmodified images (slope near zero), consistent with the results of Fig. 4. But when the responses to modified images in the ChR2 conditions were used (Fig. 8C, right), the dependence of responses on spatial contrast was largely restored with a positive slope of the regression line. Comparing the spatial contrast sensitivity values between unmodified and modified images over all cells corroborated this observation. For the *sig90/sub08* modification, for example, spatial-contrast

sensitivity values ranged approximately from zero to unity and were much larger for almost all cells as compared to unmodified images (Fig. 8D, left).

Thus, the image modifications restore sensitivity to spatial contrast and textures, despite the fact that the integration of light intensity signals by ChR2 across the receptive field is approximately linear (Fig. 1D,E and Fig. 4). The reason for this is that the image modifications include a rectification of pixel intensity values similar to commonly considered bipolar-to-ganglion cell nonlinear transmission, which avoids cancellation of activation by positive and negative contrast. Such a rectification turns high-frequency spatial contrast into a signal with an overall positive activation. In the case of the image modification, this simply occurs because the thresholding increases the average brightness of an image patch when there is spatial structure so that part of the patch is above threshold and another part below. In fact, this mechanism seems so strong in the applied modifications that the contrast sensitivity values were even larger than in the control condition (Fig. 8D, right). Thus, while we find that the image modifications do restore spatial contrast sensitivity in the ChR2 condition, a milder nonlinearity than the absolute and relatively high threshold might be more appropriate for quantitatively bringing spatial contrast sensitivity back to the natural range.

### Direct comparison of responses to modified images with control responses

The modifications applied to the natural images were mostly geared towards reducing the differences in image-response curves of RGCs under the control and the ChR2 condition. This should also make ChR2-evoked responses to individual images more similar to control responses when using modified images as compared to unmodified images. To look at this more specifically, we directly compared the average firing rate for each image and each cell between control condition and ChR2 condition with modified images, analogously to the analysis of Fig. 2C for unmodified images.

Figure 9 shows such comparison for the *sig33/sub08* modification for four sample cells, two ON and two OFF cells. The examples demonstrate that the image modifications lead to data points lying closer to the identity line, which signifies equal responses under ChR2 and control, as compared to unmodified images. For ON cells, the often apparent concave bent in the image-response curves, with stronger ChR2-evoked responses than control responses for weakly activating images, followed by plateauing ChR2-evoked responses (see Cell 1 of Fig. 9A), is smoothed out, though not completely eliminated. For OFF cells, the contrast inversion of the images alone (Fig. 9B, middle column) turns the negative correlation between ChR2 responses and control responses into a positive one, but the additional image modification further help in bringing the data points closer

**Figure 9.**
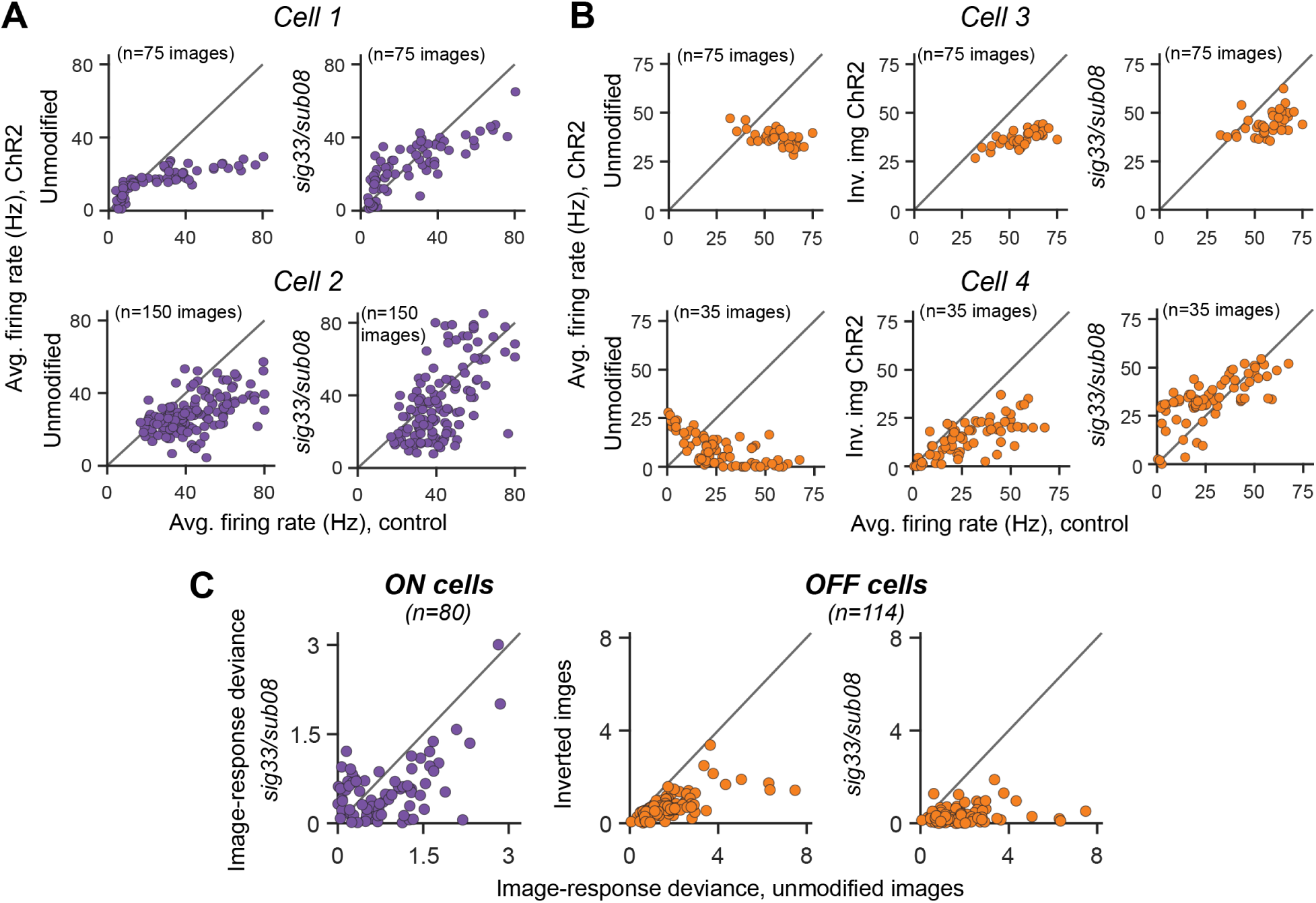
Image-by-image comparison of evoked average firing rates in the control condition versus the ChR2 condition with modified and unmodified images. Results are here shown for the *sig33/sub08* modification. **A.** Comparison of responses in the control condition versus the ChR2 condition, using unmodified images (left) or modified images (right, *sig33/sub08* modification) for two sample ON cells. **B.** Comparison of responses in the control condition versus the ChR2 condition, using unmodified images (left), images that were only contrast-inverted (middle), and fully modified images (right, *sig33/sub08* modification including contrast inversion) for two sample OFF cells. Each data point is an image. **C.** Comparison of image-response deviance in the ChR2 condition for unmodified images versus modified images (*sig33/sub08* modification) for ON cells (purple, left), versus simply inverted images for OFF cells (orange, middle), and versus fully modified images (*sig33/sub08* modification including contrast inversion) for OFF cells (orange, right).

to the diagonal (Fig. 9B, right column).

Overall, we found that the average response of the cells to the modified images in the ChR2 condition was larger than to the unmodified images and also more similar to the response level in the control condition. This is evident in the overall response ratio (ratio of average response to images in the ChR2 condition versus the control condition for each cell), which increased from 0.78 *±* 0.7 (mean *±* standard deviation, n = 125) with unmodified images to 1.21 *±* 0.91 (n = 96, *sig33/sub08* modification) for ON cells and from 0.50 *±* 0.22 (n=157) to 1.13 *±* 0.78 (n = 126, *sig33/sub08*) for OFF cells.

In order to quantify this similarity in responses between the control and the modified images in the ChR2 condition across all cells, we assessed how far the relationship between the responses in the two conditions was from the identiy line. To do so, we fitted a line passing through the origin to the data points, obtained the slope of the line, and used the absolute value of the logarithm of the slope as a measure of image-response deviance between the control condition and the ChR2 condition with a given image modification (see Methods). This image-response deviance is zero if the data points line up around the identity line, and smaller values generally indicate that image responses were more similar between control and ChR2 conditions.

Indeed, for ON cells, the image-response deviance calculated for ChR2 responses to modified images with, for example, the *sig33/sub08* modification was considerably lower (0.54 *±* 0.50, mean *±* standard deviation, n=80) than the image-response deviance obtained from ChR2-evoked responses to unmodified images (0.84 *±* 0.67, mean *±* standard deviation, *p* = 2.5 *×* 10^−5^, Wilcoxon signed-rank test, n=80, Fig. 9C, left). Similarly, for OFF cells, the image-response deviance values decreased when including image modification. The inversion of pixel contrast values alone already reduced the average image-response deviance from 1.74 *±* 1.22 (mean *±* standard deviation) for unmodified images to 0.72 *±* 0.50 (Fig. 9C, left, *p <* 10^−19^, Wilcoxon signed-rank test, n=114), and including the *sig33/sub08* modification resulted in a further decrease of the image-response deviance to 0.32 *±* 0.3 (Fig. 9C, right; *p <* 10^−15^ for comparison of *sig33/sub08* and just inverted images, Wilcoxon signed-rank test, n=114). This shows that, at least on average, ChR2-evoked responses could be made more similar to responses in control condition when image modifications were included in the stimulation.

## Discussion

The use of optogenetics for restoring vision in the case of retinal-degeneration-induced blindness has already shown promising potential in clinical trials (Sahel et al. 2021; Lam et al. 2025; Mohanty et al. 2025). However, how natural this restored vision can be and what modifications need to be made to make it more natural is still largely unknown. Here, we therefore investigated how optogenetically modified mouse retinas with ChR2 expressed in retinal ganglion cells respond to natural images. The direct comparison of photoreceptor-driven responses with ChR2-driven responses on a cell-by-cell basis revealed systematic difference in image encoding, in particular a reduction in dynamic response range, in nonlinear response components, and in sensitivity to local spatial contrast in the ChR2 condition. Image modifications that comprise spatial low-pass filtering, thresholding, and scaling of pixel intensities help reduce these differences and could become part of the stimulus encoding process from camera images to optical stimulation patterns in clinical applications of optogenetic vision restoration.

### Mechanistic differences in image encoding between optogenetic and natural stimulation

Our findings show that the changes in the encoding of natural images between photoreceptor-driven stimulation and ChR2-based activation of ganglion cells can be profound and go beyond a mere monotonic transformation of firing rates. Instead, the ganglion cells’ preferences for image features is changed so that responses can be strong with ChR2 stimulation and weak with photoreceptor stimulation for some images and vice versa for others (Fig. 2). In particular, images with fairly homogeneous spatial structure and mean light intensity near background levels could still activate RGCs under ChR2 but often fell below threshold in control conditions, whereas strongly structured images could show the opposite response relationship.

Several mechanisms appear to contribute to these differences. In particular, the approximately linear dependence of ChR2-mediated currents on light intensity over a fairly wide range (Ishizuka et al. 2006) together with the lack of retinal-circuit processing, as the ChR2 is expressed directly in the ganglion cells, supports the linear response characteristics in the ChR2 condition. Moreover, the fact that part of the light-gated current of ChR2 is non-inactivating (Ishizuka et al. 2006) leads to a persistent activation under background illumination and thus the lack of an activity threshold near this illumination level, unlike the image encoding in the control condition for most cells. Furthermore, the extension of the linear response regime towards negative Weber contrast together with the fairly linear photocurrents leads to the loss of spatial-contrast sensitivity. The latter follows because the effects of brightening and darkening within the receptive field can now cancel out, as darkening reduces ChR2 photocurrents from the corresponding receptive-field locations. Under normal stimulation, an equivalent decrease in input current to the ganglion cell is prevented for many cells by the nonlinear synaptic transmission from the presynaptic bipolar cells (Demb et al. 2001; Schwartz et al. 2012; Borghuis et al. 2013).

### Rationale of image modifications

The difference between the image-response curves in the two conditions was the primary target of our tested image modifications. Thresholding the image intensity and setting the threshold and the background to black can be viewed as the appropriate operation for reducing baseline activity and preventing activation by image parts below the original background intensity. In addition, this allowed increasing the dynamic range of the ChR2-evoked responses by scaling the pixel intensities above threshold to the entire available light-intensity range and using this full range for encoding positive activation of the cells. Alternatively, one may justify the thresholding and resetting of the background as a way to mimic processing by the retinal circuitry, which is skipped in the direct ChR2 stimulation of the ganglion cells. In particular, in normal retinal signal encoding, light adaptation to the background intensity and rectification occuring at the bipolar-to-ganglion cell synapse (Demb et al. 2001; Schwartz et al. 2012; Borghuis et al. 2013) or during spike generation induce similar thresholding operations near background intensity (Demb 2008; Jarsky et al. 2011).

Similarly, the spatial blurring was motivated by the need to avoid activation by small spots of light after introducing thresholding of pixel intensities, but also reflects spatial filtering as occurs in the retina. In fact, the two probed scales approximately reflect spatial filtering by bipolar and ganglion cell receptive fields, the main sites of excitatory signal pooling in the retina. Retinal ganglion cells in the mouse have been shown to vary in their scale of spatial integration, with some cells well approximated by linear receptive fields while others show evidence of nonlinear subunits (Karamanlis, Gollisch 2021). Thus, one might expect that different spatial scales of blurring should be appropriate for different types of ganglion cells. Although the preference for either of the two tested scales indeed varied between cells, differences in how well they worked in bringing ChR2-mediated responses closer to normal were generally rather small for our data. This suggests that the applied spatial filtering scale is not critical, as long as sufficient filtering is applied to reduce erroneous activation by contrast on very small scales.

The applied thresholding of pixel intensities is, of course, only a rough approximation of the nonlinear transformation occurring in the retina. In particular, nonlinearities of bipolar-to-ganglion cell synaptic transmission appear to be milder and more gradual (Schwartz et al. 2012). Inhibitory signals, such as cross-over inhibition from amacrine cells, may further modify signal transmission and reduce or partly linearize the threshold effect compared to what might be expected from nonlinear synaptic transmission of the excitatory signals (Werblin 2010). In this regard, image transformations with a milder nonlinearity, using perhaps incomplete rectification and some residual background light intensity might further improve the similarity between responses in the control and ChR2 condition. For higher light intensities, some saturation in the intensity transformation might also be useful to match saturation effects observed in bipolar cell input to ganglion cells (Schwartz et al. 2012).

A milder nonlinearity may also be beneficial for restoring normal ganglion-cell sensitivity to spatial contrast. While the cells displayed essentially no sensitivity to spatial structure at high spatial frequency in the ChR2 condition with unmodified images, the modifications led to an overcompensation, with spatial-contrast sensitivity even higher than in control conditions (Fig. 8C). The reason for this is that the thresholding turns any spatial-contrast signal, such as a black-white boundary, into a strong average-luminance signal, as cancellation of positive and negative Weber contrast is eliminated by the image transformation. A milder nonlinearity, with incomplete rectification, would thus bring back some of this cancellation and reduce the artificially strong spatial-contrast signal.

Optimizing the shape of the nonlinear transformation is an interesting future research direction, but might require a more cell-type-specific analysis to cope with the variability of observed image-response curves. An interesting approach could here be to compute the optimal transformation, based on an appropriate computational model of image encoding. The linear–nonlinear (LN) model (Chichilnisky 2001; Schwartz et al. 2006), for example, appears to successfully capture responses to natural images in the ChR2 condition (c.f., Fig. 7), and for the sensitivity to local spatial contrast in the control condition, the spatial contrast model (Liu et al. 2022; Sridhar et al. 2026) provides a useful extension of the LN model. Using these models together, one may hope to optimize the transformation through *in silico* approaches. Note, however, that these models typically rely on keeping basic statistics of stimulation roughly constant, such as average illumination and contrast, and that changes in stimulus statistics from the image transformations, in particular in background light level, may challenge the validity of these models.

### Cell-type-specific effects

As direct optogenetic activation of ganglion cells bypasses the retinal circuitry, much of the processing that gives the ganglion cell types their specific functional features is lost and different ganglion cells therefore resemble each other much more in their ChR2-evoked response characteristics than under normal conditions. Yet, some cell-type specificity persists in the ChR2-evoked responses to natural images, as seen by the differences in the nonlinearity of the image-response curves between OFF cells with monophasic versus biphasic temporal filters (Fig. 3C,E). Cell-type-specific differences in morphology or ionic conductances, as involved, for example, in spike generation (Margolis, Detwiler 2007; Wienbar, Schwartz 2022; Chang et al. 2024), might contribute to these differences.

More generally, however, the overall similarity of ChR2 response characteristics across cells of different functional types means that different image transformations for these types are likely required to optimally recreate their specific functional properties. This is most obvious for ON versus OFF cells, as the latter require an image inversion to match the natural contrast preference. For clinical application, stimulating ON and OFF ganglion cells independently with non-inverted and inverted images, respectively, would require selective expression of different opsins in appropriate subtypes of ganglion cells to allow independent activation through different spectral channels. To date, this remains an unsolved challenge, but the rapid development of single-cell transcriptomics (Hahn et al. 2023) and artificial-intelligence-based identification of functional promoter sequences (Avsec et al. 2021) suggest that this will be a future possibility. Meanwhile, clinical studies with optogenetic intervention at the level of ganglion cells proceed despite the unsolved problem of simultaneous stimulation of ON and OFF cells and thereby may be aided by transformations as suggested here.

ON-OFF cells provide yet another group of cells that would require independent image modifications for appropriate restoration of natural responses. In principle, this could be obtained by superposing the modified versions of the non-inverted and the inverted image, potentially with appropriate relative weighting. However, given that the primary ganglion-cell targets for clinical vision restoration, midget and parasol cells, are not of ON-OFF type, we did not investigate the effect of such transformations and instead excluded ON-OFF cells from further analyses.

Beyond the ON- versus OFF-type distinction, we aimed for generic transformations that are likely beneficial across cell types, as thresholding and spatial filtering are prevalent mechanisms of signal encoding by RGCs. The fact that the selected image modifications worked well for most ganglion cells (though not for all), independent of the specific type, also suggests that the effect is robust against variations in response characteristics. This also lets us expect that the findings do not strongly depend on variations in opsin-expression level in the ganglion cells, which are likely higher when viral transduction is used to express the opsin in ganglion cells as compared to the transgenic mice used here. Yet, future studies with viral-vector-mediated opsin expression in wild-type mice should test this directly. Similarly, it seems likely that the usefulness of such transformations generalizes to light-activated ion channels other than ChR2, as similar response characteristics can be expected for many optogenetically applied opsins.

### Alternative optogenetic interventions and future directions

Ganglion cells are considered to be the most resilient retinal nerve cells in degenerative conditions such as retinitis pigmentosa (Mazzoni et al. 2008; Jones et al. 2016; Pfeiffer et al. 2020). Furthermore, they are likely the physically easiest target for optogenetic interventions in the retina (Lindner et al. 2022; Prosseda et al. 2022) because delivery by intravitreal injections let the opsin-carrying viral vectors directly encounter the ganglion cells in the retina and because subretinal injections, which may be chosen to target bipolar cells, are more difficult and risky in terms of physical damage.

Yet, targeting cells earlier in the retinal processing pathway than ganglion cells remains a fascinating alternative approach, in particular in cases where degeneration and remodeling has not yet strongly impacted the inner nuclear layer (Jones et al. 2016; Pfeiffer et al. 2020). For example, expression of opsins in bipolar cells (Lagali et al. 2008; Doroudchi et al. 2011; Gaub et al. 2015; Macé et al. 2015; Kralik et al. 2022; Rodgers et al. 2025) but also in remaining cone cell bodies (Busskamp et al. 2010) or amacrine cells (Khabou et al. 2023) currently makes differentially activating ON and OFF cells easier than expression in ganglion cells. Furthermore, retaining more of the processing by the retinal circuit may lead to more natural responses even without image modifications and can help preserve response diversity and specific functionality (Rodgers et al. 2023; Rodgers et al. 2025). Yet, it seems likely that appropriate modifications will still be helpful in the case of targeting bipolar cells, as retinal processing is still partly bypassed. In what way the appropriate stimulus modifications differ when ganglion cells or bipolar cells are targeted and how the inclusion of such modifications affects the ability to create responses as normal as possible by optogenetic bipolar-cell activation remain important open questions.

While we applied spatial low-pass filtering by blurring the images to reduce responses by small stimulus components, another standard spatial-filtering operation of the retina is a certain level of high-pass filtering induced by the center–surround structure of receptive fields. For the present data, receptive-field estimates did not show a strong surround component, as is often the case for receptive fields obtained under stimulation with spatiotemporal white noise (Kerschensteiner et al. 2008; Koehler et al. 2011; Cowan et al. 2016; Wienbar, Schwartz 2022). Furthermore, the image-response curves based on these estimated receptive fields appeared to capture the overall response structure well. Thus, the lack of such high-pass filtering in the ChR2 condition does not seem a major source of error. Yet, adding such a filtering component in the future may help bring natural and optogenetically evoked image responses closer together, for example, by reducing some of the overshooting responses to modified images in the ChR2 condition, which could reflect a lack of surround-mediated suppression. In this respect, it is noteworthy that past work created center–surround receptive fields with optogenetic activation of ganglion cells by expressing different opsins differentially in the soma and dendrites of the cells (Greenberg et al. 2011), though obtaining natural spatial scales then also required appropriate blurring of the images.

Finally, stimulus modifications with spatial band-pass filtering could be combined with appropriate temporal filtering, while retaining the nonlinear transformation and scaling of pixel intensities. The temporal transformation should include a low-pass filter to counteract the faster response kinetics under ChR2 stimulation. The added temporal filtering would then make the stimulus modifications applicable to natural movies and thereby allow assessing how well ChR2-evoked responses with simple stimulus transformations could mimic natural responses in a real-world scenario.

## Methods

### Experimental design and statistical analysis

Mice with Cre-dependent Channelrhodopsin-2 (Ai32(RCL-ChR2(H134R)/EYFP) strain; Jackson Laboratory, stock no. 024109) and mice with Cre expression in glutamatergic neurons (Vglut2-ires-cre knock-in (C57BL/6J) strain; Jackson Laboratory, stock no. 028863) were purchased from Charles River and mated to obtain offspring that expressed Channelrhodopsin-2 (ChR2) in retinal ganglion cells (RGCs). Retinas from these transgenic mice of either sex (5 male, 5 female), aged four to ten months, were used for experiments. All mice were housed in a 12 h light/dark cycle. Experimental procedures were in accordance with institutional and national guidelines and approved by the institutional animal care committee of the University Medical Center Göttingen (protocol number T 23.26). No statistical methods were used to predetermine sample size. For statistical comparisons of the characteristics of the recorded cells between different stimulation conditions, we applied a two-sided Wilcoxon signed-rank test. All statistical procedures were conducted using MATLAB R2019b functions.

### Tissue preparation and electrophysiology

Mice were dark-adapted for approximately two hours and sacrificed by cervical dislocation. The eyes were removed and dissected in Ames’ medium (Sigma Aldrich), supplemented with 4 mM D-glucose monohydrate (Carl Roth) as well as 20 mM sodium bicarbonate (Merck Millipore) and oxygenated (95% O_2_/5% CO_2_) to maintain a pH of 7.4. The cornea, lens, and vitreous humor were carefully removed, and the eyecups were cut into halves to allow separate recordings with each half. The tissue preparation was performed under infrared illumination, using a stereomicroscope equipped with night-vision goggles.

To record ganglion-cell spiking activity, retina pieces were isolated from the eyecup and placed ganglion cell-side-down on planar multielectrode arrays (MultiChannel Systems, 252 electrodes, electrode diameter 8 or 10 µm, minimal electrode distance 30 or 60 µm), which were coated with poly-D-lysine (1 mg/mL; Millipore). Throughout the recording, the retinal tissue was perfused with pre-heated (33°C), oxygenated Ames’ medium at a rate of 4–5 mL/min. The bath solution and the recording chamber were maintained at 32°C-33°C using an inline heater and a heating element below the array.

For the parts of the recording that aimed at investigating responses under activation of ChR2, a mixture of pharmacological drugs was applied together with the Ames’ medium to the same piece of retina in order to suppress any photoreceptor-mediated neuronal activity. In particular, we used CNQX (80 µM, CNQX disodium salt, Abcam and NeoBiotech) to block glutamate receptors in ganglion cells, DL-AP4 (40 µM, Biotrend) to block metabotropic glutamate receptors in ON bipolar cells, and ACET (1 µM, Tocris) to block kainate glutamate receptors in OFF bipolar cells. Recordings were resumed 7-10 min after starting the drug application, which continued till the end of the experiment. For each experiment, the pharmacological solutions were freshly prepared from stock solutions.

The recorded voltage traces were amplified, bandpass-filtered (300 Hz to 5 kHz), and sampled at 25 kHz for digital storage. To extract spike times, we applied a modified version of the spike-sorting software Kilosort (Pachitariu et al. 2016; Pachitariu et al. 2024), modified code available at https://github.com/dimokaramanlis/KiloSortMEA), and manually curated the spike clusters with the Phy2 (https://github.com/cortex-lab/phy) software package. For further analysis, we used only well-isolated clusters with minimal refractory-period violations.

### Visual stimulation

Custom-made software based on Visual C++ and OpenGL was used to generate visual stimuli. These were displayed on a gamma-corrected DLP projector (E4500MKII, EKB technologies, equipped with UV, blue, and green LEDs) with 60 Hz refresh rate and a resolution of 1280 *×* 800 pixels and projected onto the retina with a custom-built lens system (pixel size 5.6 µm *×* 5.6 µm or 6.2 µm *×* 6.2 µm on the retina, depending on which of two available experimental setups was used). Before the start of the experiment, the stimulus display was focused on the photoreceptor layer by visual monitoring via a light microscope.

For recording photoreceptor-mediated responses (also called “control condition” in the following), overall light intensity was reduced by neutral density filters placed in the projection path, and all stimuli had a mean light intensity on the retina of 6.7 mW/m^2^ in the low photopic range, using all three LEDs (UV, blue, and green). This light intensity is several orders of magnitude below the typical intensities needed for activating ChR2 (Lin 2011) so that potential contributions from ChR2 to the activation of the retina are negligible in this condition. The isomerization rates for the different types of photoreceptors were calculated to be (in isomerizations per photoreceptor and second) around 4000 for rods, 250 for S-cones, and 1600 for M-cones, based on collecting areas and spectral sensitivities from https://github.com/eulerlab/open-visual-stimulator/blob/master/ `calibration_mouse/stimulator_calibration.ipynb` and (Field, Rieke 2002; Nikonov et al. 2006). For stimulation of ChR2 (also called “ChR2 condition”), neutral density filters were partly removed and only the blue LED (peak wavelength 460 nm) of the projection system was used, yielding a mean irradiance for all stimuli of 0.7 mW/mm^2^ on the retina. This light level was chosen based on the light intensities used in previous studies to activate ChR2 (Nagel et al. 2005; Lin 2011; Britt et al. 2012) and ensured that the range of displayed intensities fell into the sensitivity range of ChR2 while avoiding saturation (Lin 2011).

### Responses to light-intensity steps

To characterize preferred contrast polarity and response kinetics of the recorded cells, we analyzed their responses to the ‘chirp’ stimulus as used by Baden et al. (2016). The stimulus consists of three parts: step-like changes in light intensity, frequency sweep, and contrast sweep. For our analysis, we only focused on step-like changes in light intensity, comprising a sequence of 4 s of darkness (*−*100% Weber contrast), 3 s of high light intensity (+100%), 3 s of darkness (*−*100%), and 2 s of background illumination (0%). The chirp stimulus was repeated for 15 trials, and spikes were binned at 10 ms bin size to compute peri-stimulus time histograms (PSTHs).

### Characterizing temporal filters and receptive fields

To estimate the receptive field (RF) of each RGC, a spatiotemporal binary white-noise stimulus (100% Michelson contrast) with a checkerboard layout was used. It consisted of flickering squares (each 7 *×* 7 projector pixels and around 40 µm to the side), updated independently and randomly at the projector refresh rate (60 Hz). We measured the receptive field of a cell by first calculating the spatiotemporal spike-triggered average (STA) from the cell’s recorded spikes (Chichilnisky 2001). We separated the spatial and temporal components of the STA by selecting the stimulus square with the absolute maximum value in the STA after smoothing with a spatial Gaussian filter of 20 µm standard deviation. The time course (before spatial smoothing) of this selected square was taken as the temporal filter of the cell. To reduce noise, we restricted subsequent computations to a fixed number of pixels around the selected square, cropping the STA to a square region of 700 µm per side.

To obtain an estimate of the RF in pixel-wise fashion, we computed for each pixel the dot product of the pixel’s temporal sequence in the STA with the temporal filter. For plotting RFs of OFF cells in the control condition, we adjusted the sign of the spatial filter to show a negative peak. To assess the RF size, we fitted a two-dimensional Gaussian to the RF estimate and defined the size as the diameter of a circle with equal area as inside the 1.5-*σ* elliptical contour of the fit. The RF outlines displayed in the figures mark the 1.5-*σ* contours of the fit. To obtain the time-to-peak in the temporal filter, we fitted a second-degree polynomial around the primary peak (positive peak for ON-type filters, trough in case of OFF cells in the control condition, using five data points around the one with the highest absolute value) and took the corresponding time of the maximum (or minimum in case of OFF-type filters) as the time-to-peak. Both the RF estimate and temporal filter were normalized to unit Euclidean norm.

### Assessing spatial nonlinearities with contrast-reversing gratings

Spatial integration of the RGCs was characterized by analyzing responses to contrast-reversing gratings as previously described (Karamanlis, Gollisch 2021). The retina was stimulated with full-field square-wave gratings of 100% Michelson contrast, and contrast was reversed every one second. The gratings were presented in order from higher to lower spatial frequencies, applying seven spatial frequencies with bar widths ranging from 25 µm to 800 µm and sampling two to eight equidistant spatial phases for some of them (e.g., two for 50, four for 200, and eight for 400 µm bar width). Each grating at a given spatial frequency and phase was shown for 20 reversals. Between the presentations of gratings, a gray screen at background intensity was displayed for 1.2 s. PSTHs were calculated over one reversal period for each spatial frequency and phase with 10 ms bins.

We obtained a nonlinearity index in response to reversing gratings as described previously (Karamanlis, Gollisch 2021), based on the index calculation in (Hochstein, Shapley 1976). In brief, we computed the discrete temporal Fourier transform of the PSTH for each combination of spatial frequency and phase. We then extracted the maximum amplitude *F* 1 at the contrast-reversal frequency across all spatial frequencies and phases as well as the maximum amplitude *F* 2 at twice the reversal frequency across all PSTHs with high spatial frequencies (bar width *≤* 200 µm). The assessment of *F* 2 was here restricted to high spatial frequencies to avoid spurious high values from non-sinusoidal temporal response profiles at low spatial frequency, which did not reflect the typical frequency doubling of nonlinear spatial integration. The nonlinearity index was calculated as the ratio *F* 2*/F* 1. A large nonlinearity index value (typically *≥* 1) indicates nonlinear spatial integration, and a low value (e.g., close to zero) indicates linear spatial integration by the analyzed cell.

### Responses to natural images

We selected and displayed natural images as previously described (Karamanlis, Gollisch 2021). Briefly, images were selected from the van Hateren Natural Image Dataset (van Hateren, van der Schaaf 1998), the McGill Calibrated Colour Image Database (Olmos, Kingdom 2004), and the Berkeley Segmentation Dataset (Arbeláez et al. 2011), converted to grayscale if originally given as color images, and resized to 512 x 512 pixels by cropping (and upsampling if needed). Pixel intensities of all images were scaled so that the mean intensity of all images was equal to background. Occasional pixel intensity values larger than twice the background (maximum projection intensity) were then clipped at this value. The images were encoded at 8-bit color depth, presented centered on the multielectrode array, and covered a region of 2.9 mm *×* 2.9 mm or 3.2 mm *×* 3.2 mm on the retina, surrounded by background intensity.

We also included contrast-inverted images (see “Modifying natural images” below) for the purpose of comparing responses of OFF cells between the control and the ChR2 condition. The inverted images were generally treated as separate, independent images (including when analyzing ON cells), except for the specific analyses of OFF cells when image inversion was included in the image modifications. In that case, each image from the control condition was paired with its inverted (and potentially further modified) version in the ChR2 condition for comparison.

The images (300, 220, 150, or 70 images from the combined databases, including the inverted versions, with exact number depending on the experiment) were presented individually for 200 ms each, separated by 600 ms of background illumination, and the sequence of images was repeated for 10 trials, each time with a different pseudo-random permutation of the order of images. For each cell, firing-rate profiles for each image were calculated as PSTHs with 10 ms bins over a time window that also spanned some time before and after the image flash. The average firing rate for each image was taken as the mean firing rate during the 200 ms image flash.

### Modifying natural images

In addition to presenting the original natural images to the retina as described above, we also applied various modifications to the images for displaying them to the retina in the ChR2 condition.

### Inverted images

Images were inverted by flipping the sign of the Weber contrast value for each pixel.

#### Thresholded images

Pixel intensities were thresholded at the background light intensity by subtracting the background value and setting resulting negative values to zero. Subsequently, all pixel intensity values were scaled to recover the original range of available intensity values, here obtained by multiplying all pixel values by a factor of two. These images were presented in the same way as the original images, but on a black background (corresponding to an intensity value of zero in the images) and separated by the same black background.

### Blurred and thresholded images

Further image modifications included spatial filtering (blurring) and different levels of thresholding. The images were blurred by convolution with a two-dimensional spherically symmetrical Gaussian filter with standard deviation *σ* of either 33 µm or 90 µm to test different levels of blurring. After spatial filtering, we again thresholded and scaled the intensity values, as explained above (“Thresholded images”), though here we subtracted not only the background intensity, but also an additional subtraction level *M* of either 8% or 16% of the maximum available intensity. The images were then again scaled to restore the maximum projection intensity. Image sets modified by the above parameters are denoted by the sigma value of spatial filtering and the subtracting level *M*. For example, images modified by spatial filtering using *σ* = 33 µm and *M* = 8% are referred to as *sig33/sub08*, and images modified by spatial filtering using *σ* = 90 µm and *M* = 16% are referred to as *sig90/sub16*.

### Image-response curves for natural images

To analyze how the activation of a cell’s receptive field relates to the evoked response under natural images, we obtained image-response curves in the following way. For each cell and image, we calculated the mean stimulus intensity inside the receptive field *I_mean_* (Liu et al. 2022; Sridhar et al. 2026) by weighting the Weber contrast values of the image pixels with the values of the RF estimate at the corresponding locations and then averaging over the cropped region of the spatiotemporal STA (see section on “Characterizing receptive fields” above). The corresponding response was computed as the average firing rate during the time of the image presentation and over trials. The image-response curve of each cell is obtained by plotting the average firing rates against the corresponding *I_mean_*values. In all figures, the *I_mean_*values were multiplied by 1,000 for easier readability of the plots.

The *I_mean_* values for each cell and image were calculated separately for the control and the ChR2 condition by using the corresponding RF estimates from responses to white-noise stimuli in each condition. The cells that had a rightward-rising image-response curve in the control condition (i.e., increasing firing rate with increasing *I_mean_*) were classified as ON cells, and the cells with leftward-rising image-response curves in the control condition (i.e., increasing firing rate with decreasing *I_mean_*) were classified as OFF cells. Cells that could not be clearly classified as either ON- or OFF-type (either because the image-response curves were too noisy or unstructured or because they were U-shaped with increasing responses in both positive and negative directions from the middle, as characteristic for ON-OFF cells) were excluded from further analysis (see “Cell exclusion” below).

When computing image-response curves in the ChR2 condition in response to modified natural images, we used the average firing rates as obtained for the modified images but kept the *I_mean_*values as computed from the corresponding unmodified images in the ChR2 condition. The reasoning behind this is that the unmodified image is what needs to be encoded and the applied image manipulations are considered part of the encoding process, together with the ChR2-driven activation of the RGC.

### Nonlinearity index of image-response curves

To quantify the degree of nonlinearity of each image-response curve, we computed a nonlinearity index by first fitting two straight lines to the data points in the curve, one for all points with *I_mean_ <*0, yielding the slope value *S*_1_, and one for all points with *I_mean_ >*0, yielding the slope value *S*_2_. The ratio between the difference and the sum of the two slope values, (*S*_2_ *− S*_1_)*/*(*S*_2_ + *S*_1_), was taken as the nonlinearity index. For OFF cells in the control condition, the value of the index was multiplied by *−*1 to reflect the leftward-rising shape of the curve. The index typically takes values between unity (highly nonlinear, rectified response) and zero (highly linear response). Any values larger than unity and lower than zero were clipped to unity and zero, respectively.

### Comparing image-response curves between conditions

For comparing the image-response curves across cells and different conditions, we used a least-squares criterion to fit the data with a parametrized softplus function *y* = *A* ln(1+*e^Bx^*^+^*^C^*)+*D*, where *A* is the scaling factor along the *y*-axis (firing rate), *B* is the scaling factor along the *x*-axis (*I_mean_*), *C* sets the horizontal shift of the curve, and *D* corresponds to the level of background activity. The initial values used for fitting the four parameters were the maximum average firing rate over all images for *A*, 10^3^ for *B*, zero for *C*, and the baseline firing rate for *D*. The latter was obtained as the level of spontaneous activity of the cell, calculated as the average firing rate over the 200 ms periods prior to the image presentations. For OFF cells under control conditions, a negative sign was included in the exponential function before *B* to reflect the increasing response *y* for decreasing *x* while keeping *B* positive.

The fitted softplus functions were then used to quantify how well the image-response curves in the ChR2 condition for different modifications reproduced the image-response curve of the control condition. We did so by computing a distance measure between the softplus functions from the control condition and from the ChR2 condition with a given image modification scheme by averaging the absolute values of the difference of the two curves over all points with *I_mean_* values corresponding to data points of the control condition.

### Spatial contrast sensitivity for natural images

To assess the influence of spatial image structure beyond the mean stimulus intensity inside the receptive field, we computed the local spatial contrast sensitivity (SC sensitivity) of each cell as previously described (Karamanlis, Gollisch 2021). Briefly, for each cell and each image, the local spatial contrast (SC) of an image was calculated as 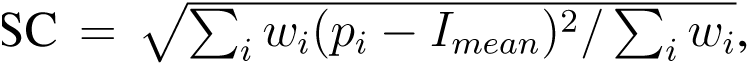 where *i* runs over all image pixels inside the cropped region of the STA of a cell, *p_i_* is the pixel value (Weber contrast) of the image, *w_i_* is the pixel weight as given by the RF estimate, and *I_mean_* is the mean stimulus intensity as described above. All values smaller than zero in *w_i_* were clipped to zero to avoid negative contributions to the SC computation. We then sorted all images according to their *I_mean_* values and formed exclusive pairs from neighboring images in the sorted list, so that images with similar *I_mean_*values were paired. For each pair, we computed the differences between SC values and corresponding firing rates (normalized by the maximum firing rate across all images), respectively. Finally, the SC sensitivity of a cell was defined as the slope of a fitted linear regression line, describing the dependence of firing-rate differences on SC differences.

### Direct comparison of responses between conditions

To directly compare responses to individual images in control condition and in one of the different ChR2-condition scenarios, the average firing rate during the image flash for each image in control condition was plotted against the average firing rate of the same image (potentially including image modifications) in the ChR2 condition for each cell. We then fitted a straight line, passing through the origin, to these data points and calculated the absolute values of the natural logarithm of the slope. This value is referred to as the image-response deviance. The smaller the image-response deviance (i.e., the closer to zero), the higher the similarity between the responses in the two conditions. The logarithm was included in the calculation of the image-response deviance because a slope of unity then yields a deviance of zero, reflecting the best possible match between control and ChR2 responses, and because, for example, slope values of two and one half correspond to equally strong deviations from this best possible match. For this analysis, two OFF cells were excluded because of their poor responses to unmodified images in the ChR2 condition, which resulted in a slope of zero of the fitted line. Furthermore, one recording was excluded from this analysis because of a slight lateral shift of the projected images between control and ChR2 conditions.

### Cell exclusion

We excluded cells for which no clear receptive field was obtained from the spike-triggered-average analysis under white-noise stimulation in either control or ChR2 condition. This was assessed by whether an unambiguous receptive-field location was visible in the spatial component of the spike-triggered average and whether this was well-fitted by a two-dimensional Gaussian function. A lack of a clear receptive field typically corresponded to small firing rates, below about 5 Hz, in response to the white-noise stimulus.

We also excluded cells lacking a clear image-response curve that could be classified as either ON-type (rising) or OFF-type (falling) in the control condition. This included cells where the image-response curve was essentially unstructured as well as cells where the image-response curve was U-shaped, rising to similar levels for large positive and strongly negative *I_mean_* values, which corresponds to the case of an ON-OFF cell. Finally, cells that overall did not respond well to the natural images in either the control or the ChR2 condition (as assessed by maximum firing rates averaged over the presentation of individual images below about 20 Hz) were excluded. In total, 355 cells were excluded by the criteria on the image-response curves, whereas 125 ON cells and 157 OFF cells were included for final analyses.

### Immunohistochemistry

To check expression of ChR2, we performed immunohistochemistry on retinas of six transgenic mice of either sex (2 male, 4 female), aged between four and twelve months. Retinas of two of these mice were also used for ganglion-cell recordings, the other mice were littermates of the ones used for ganglion-cell recordings. The retinas were isolated as done for electrophysiology and fixed in 4% para-formaldehyde (36-38% formaldehyde in water, Sigma-Aldrich, dissolved in 10 mM PBS, Sigma-Aldrich) for 30 min. Subsequently, the retinas were washed in 10 mM PBS (3 washes, 10 min each) and incubated in blocking solution for 2 hours at room temperature. The blocking solution consisted of 3% donkey serum (Sigma-Aldrich, D9663) in 10 mM PBS with 1% TritonX-100 (Sigma-Aldrich, T8787). This was followed by incubation in primary antibody solution at 4°C on a shaker for 7 days.

The retinas were co-stained for RBPMS (RNA-binding protein with multiple splicing, which is expressed in RGCs) with anti-RBMPS antibody (raised in guinea pig, Merck, LOT#3282144) and for EYFP (enhanced yellow fluorescent protein, which is a reporter gene in the studied transgenic mice for ChR2) using anti-GFP antibody (raised in rabbit IgG, ThermoFischer, Cat#A11122). Both primary antibodies were applied in a 1:500 dilution in the blocking solution. After 7 days, the retinas were washed in 10 mM PBS (5 washes, 15 min each) and incubated in secondary antibody solution at room temperature for two hours. AlexaFluor 647 (Dianova, anti-guinea pig, diluted in glycerol 1:2) and AlexaFluor 488 (Invitrogen, LOT #2156521, anti-rabbit IgG) were used in a 1:500 dilution in the blocking solution as secondary antibodies. The retinas were then washed in 10 mM PBS and mounted on a slide with Aqua Polymount (Polysciences Inc, LOT #A815745). Images of the retinas were obtained with a Leica SP8 DMi8 confocal microscope, using either 63x or 20x objective magnification, and processed using Image J.

## Data availability

The spike train data analyzed here have been made publicly available at https://gin.g-node.org/gollischlab/Ramakrishna_Gollisch_2026_Mouse_RGC_Spiking_ChR2_Natural_Images (DOI: 10.12751/g-node.h9gnd4).

## Acknowledgments

This work was supported by the Deutsche Forschungsgemeinschaft (DFG, German Research Foundation) – project numbers 528760423 (SFB 1690/1, “Disease Mechanisms and Functional Restoration of Sensory and Motor Systems”, project B05 to T.G.) and 515774656 (to T.G.) – and by the Else Kröner Fresenius Foundation through the Else Kröner Fresenius Center for Optogenetic Therapies. We would like to thank D. Karamanlis, M. Khani, and M. Weick for their help in setting up of the stimulus-projection system and in preparing visual stimuli and analysis pipelines.

## Author contributions

V.R. and T.G. conceived and designed experiments and data analysis methods. V.R. performed experiments, analyzed data, and prepared figures. V.R. and T.G. wrote the paper.

